# Rewiring of the three-dimensional genome encodes regenerative potential in the adult central nervous system

**DOI:** 10.64898/2026.03.12.711224

**Authors:** Anisha S. Menon, Manojkumar Kumaran, Faheem Farooq, Deepta S Beji, Dhruva Kumar Kesireddy, Yogesh Sahu, Ishwariya Venkatesh

**Affiliations:** Council of Scientific and Industrial Research (CSIR) – Centre for Cellular and Molecular Biology (CCMB), Hyderabad 500007, Telangana, India; Academy of Scientific and Innovative Research (AcSIR), Ghaziabad 201002, Uttar Pradesh, India

## Abstract

The failure of adult central nervous system neurons to regenerate after injury has been attributed to transcriptional and epigenetic barriers, but whether three-dimensional genome organization constitutes an independent regulatory layer encoding regenerative potential remains unknown. Here we present the first genome-wide map of chromatin compartments, topologically associating domains, and loops across postnatal development, adult homeostasis, and spinal cord injury in the mouse motor cortex. Postnatal maturation progressively consolidates a growth-restrictive three-dimensional architecture, and spinal cord injury alone partially reverses this consolidation, re-engaging neonatal gene programs through reorganized but functionally recapitulative architecture despite minimal transcriptional activation. This reversion is directed rather than stochastic, preferentially targeting pro-growth gene networks, and reveals a latent three-dimensional memory of developmental growth states in the adult cortical genome. Strikingly, NR2F6, a transcription factor that promotes corticospinal axon regeneration, extends this reversion beyond the neonatal state toward an earlier embryonic chromatin configuration, a depth of developmental plasticity that injury alone cannot reach. These findings establish three-dimensional genome topology as a regulatory layer encoding regenerative potential in adult cortical neurons, demonstrating that successful CNS regeneration requires accessing embryonic rather than merely neonatal chromatin states, and reframing regenerative failure as a topological problem with new therapeutic targets.

## Introduction

The adult mammalian central nervous system (CNS) fails to regenerate axons after injury, with devastating consequences for spinal cord injury and stroke^1–5^. Unlike peripheral neurons, which activate a robust transcriptional regeneration program following axotomy^6^, corticospinal neurons remain largely transcriptionally silent. Two major barriers have been identified: an inhibitory extracellular environment created by myelin-associated molecules and the glial scar^7–10^, and an intrinsic failure to reactivate developmental growth programs. While transcription factor overexpression^11–17^, epigenetic remodeling^18–22^, and mTOR activation^23,24^ can induce regeneration,even successful interventions produce limited axonal growth, suggesting deeper regulatory constraints remain.

These constraints are embedded in the chromatin landscape. Pro-growth loci remain inaccessible following injury^18,25^, restricting transcription factors such as KLF6 and SOX11 to already-accessible genes^11,12,16^ and developmental growth loci progressively accumulate repressive marks during postnatal maturation^18,26–29^ suggesting regenerative failure reflects a developmental restriction rather than a purely injury-induced state. Chromatin remodeling factors including PATZ1 can partially reverse these changes^25,30^, and our prior work showed that NR2F6 occupies a large fraction of growth-associated enhancers in injured neurons^13^. However, accessibility and histone modifications describe only the local regulatory state of individual loci, and do not capture how those loci are spatially organized within the nucleus, even though spatial proximity determines whether enhancers can contact their target promoters^31–34^. The incomplete transcriptional response to injury, in which chromatin becomes more accessible at growth-associated loci without robust gene activation, suggests a structural barrier beyond the nucleosomal level.

Three-dimensional (3D) genome organization provides this missing regulatory layer^35–39^. Chromatin is hierarchically organized into megabase-scale A/B compartments^32,40,41^, topologically associating domains (TADs) maintained by CTCF and cohesin^42–47^, and focal enhancer-promoter loops that provide the most immediate structural basis for transcriptional activation^32,42,48–50^. Many regeneration-associated enhancers lie at considerable distances from their target genes, implying dependence on long-range looping that may be disrupted in the adult. Critically, the transcriptional silencing of growth-associated genes during postnatal maturation coincides with TAD boundary reinforcement and compartment segregation strengthening, raising the possibility that architectural consolidation is causally linked to transcriptional restriction and that the loss of regenerative competence may be encoded within 3D genome topology.

Despite the central role of 3D genome architecture in gene regulation, its contribution to CNS regeneration remains unexplored. Existing studies have focused on neuronal specification during embryonic development^51–53^ or enhancer-promoter looping in peripheral neurons^50^, leaving the architectural transitions associated with the loss of growth competence and their potential reactivation after injury unknown. This gap makes it difficult to explain why transcription factor interventions sometimes fail to activate regenerative gene programs, or why chromatin accessibility gains after injury do not reliably translate into transcriptional activation.

Here we generate genome-wide Hi-C maps of chromatin compartments, TADs, and loops in mouse motor cortex across postnatal day 0 (P0), uninjured adult, and spinal cord-injured states, integrated with NR2F6 overexpression, to ask whether successful regeneration is associated with 3D genome reversion toward developmental states. We report three principal findings. First, postnatal maturation drives large-scale compartment reorganization, with 15.6% of the genome switching compartment identity as neurons transition from growth-permissive to growth-restricted states, accompanied by TAD boundary reinforcement and loop stabilization at pro-growth loci. Second, spinal cord injury partially reverses these transitions in a directed rather than stochastic manner, selectively dismantling boundaries strengthened during maturation and re-engaging neonatal gene programs through reorganized but functionally recapitulative architecture, revealing that adult cortical neurons retain a latent architectural memory of their neonatal growth state. Third, NR2F6 does not simply amplify the injury response but pushes chromatin architecture beyond the neonatal configuration toward an earlier embryonic state, engaging non-overlapping loci with a stronger re-activation bias and superior loop formation. Together, these findings establish 3D genome topology as an independent regulatory layer encoding regenerative potential in adult cortical neurons and identify developmental chromatin reversion as a hallmark of successful CNS regeneration

## Results

### Establishing a high-resolution map of three-dimensional genome organization across motor cortex states

To map chromatin topology relevant to regenerative capacity, we selected postnatal day 0 and adult motor cortex as primary comparison points. By P0, cortical neurons have exited the cell cycle and established their molecular identity yet retain active axonal growth, representing a uniquely informative window in which neuronal differentiation is complete but intrinsic growth programs remain accessible. Comparing P0 to adult cortex captures the architectural transition from growth-permissive to growth-restricted chromatin state.

We performed in situ Hi-C on motor cortex tissue from P0, uninjured adult, and adult mice at 7 days following thoracic spinal cord injury. Libraries were generated through chromatin crosslinking, restriction digestion, proximity ligation, and high-throughput sequencing, followed by standardized computational pipeline for alignment, quality control, compartment calling, TAD detection, and loop identification (**Figure 1A,B**). Distance-dependent contact probability curves displayed highly similar decay profiles across all three conditions, confirming comparable polymer folding properties and library quality (**Figure 1C**). Principal component analysis of normalized contact matrices revealed robust A/B compartment segregation across all conditions, with PC1 tracks along representative chromosomes showing highly conserved large-scale compartment patterns between P0, adult, and injured states (**Figure 1D**). Insulation scores calculated at 50 kb resolution confirmed that TAD boundary positions were reproducibly detected and largely conserved across conditions (**Figure 1E**), providing a robust framework for systematic analysis of condition-specific chromatin reorganization.

**Figure 1:**
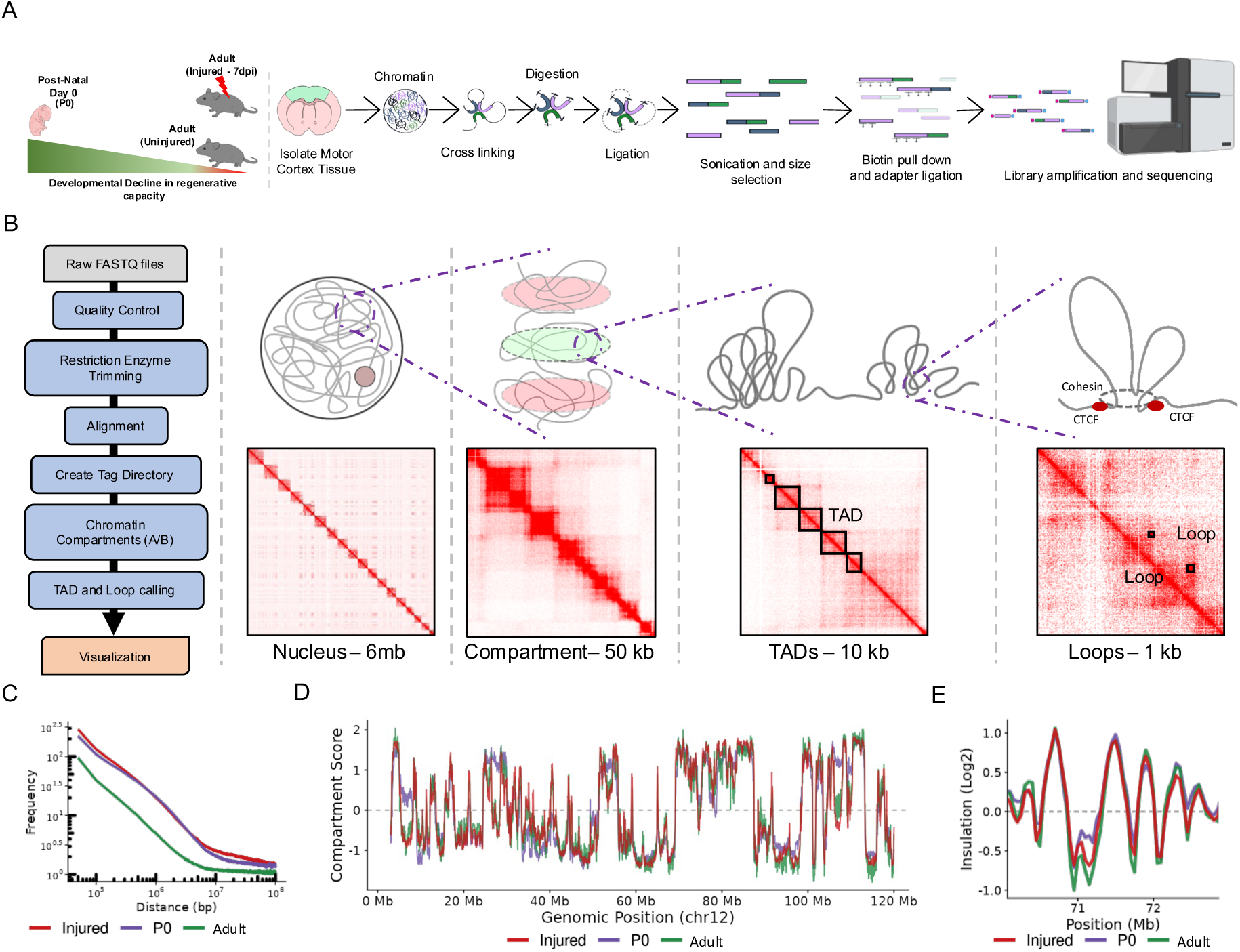
Establishing a high-resolution map of three-dimensional genome organization across motor cortex states. (A) Hi-C experimental workflow. Schematic representation of the Hi-C protocol from tissue and cell collection through chromatin extraction, crosslinking, restriction enzyme digestion, proximity ligation, DNA purification, library preparation, and high-throughput sequencing. (B) Bioinformatics pipeline and chromatin organization visualization. Left panel shows the computational workflow from raw FASTQ files through quality control, restriction enzyme trimming, alignment, tag directory creation, chromatin compartment (A/B) identification, TAD and loop calling, to final visualization. Right panel displays representative Hi-C contact maps from P0 samples illustrating hierarchical chromatin organization at multiple scales: chromatin compartments (A/B), topologically associating domains (TADs), and chromatin loops. (C) Distance-Dependent Contact Decay. Log-log plot displaying the average contact frequency (y-axis) as a function of genomic distance in base pairs (x-axis) at 50 kb resolution. The similar slopes across P0 (purple), Adult (green), and Injured (red) samples indicate consistent polymer folding properties and comparable library quality, despite differences in sequencing depth (D) A/B Compartment Analysis. First principal component (PC1) eigenvectors plotted along the entire Chromosome 12. Positive values correspond to the active A-compartment, while negative values represent the inactive B-compartment. The dashed line at y=0 marks the compartment transition. Global compartmentalization patterns are highly conserved across all three conditions (E) Insulation Score Profiles. Insulation scores (Log2) calculated at 50 kb resolution for a representative 3 Mb region (70 to73 Mb) on Chromosome 12. Local minima (dips) in the signal correspond to Topologically Associating Domain (TAD) boundaries / CTCF-enriched regions.

### Postnatal maturation reshapes chromatin compartment organization and attenuates global compartment segregation

We next asked how large-scale genome organization changes as neurons transition from growth-permissive postnatal to growth-restricted adult states. Approximately 15.6% of the genome underwent compartment switching during maturation, with 8.7% shifting from A to B (n=8,847 bins) and 6.9% from B to A (n=6,956 bins) against stable fractions of BB (n=46,285, 45.6%) and AA (n=39,401, 38.8%) (**Figure 2B, Supplementary Table S2**). Multi-resolution Hi-C contact maps revealed visibly sharper plaid interaction patterns in the adult relative to P0, suggesting refinement of compartment boundaries with maturation (**Figure 2C**). Paradoxically, saddle plot analysis revealed that homotypic interactions were stronger at P0, with a net reduction in compartment segregation strength in the adult, reflected by a delta-S of -0.89 (**Figure 2D,E**). Global compartment strength decreased markedly from 1.534 at P0 to 0.643 in the adult (Figure 2F), and the compartment enrichment ratio shifted from -0.056 at P0 to 0.104 in the adult, indicating redistribution from B-compartment-dominated interactions at P0 toward relatively enriched A-compartment homotypic contacts in the adult (**Figure 2G**). Gene ontology analysis of switching regions revealed functional stratification: regions gaining active compartment status were enriched for synaptic transmission, axon guidance, and neuronal differentiation, while regions shifting to B compartment were enriched for early developmental, proliferative, and innate immune signaling processes, reflecting progressive silencing of neonatal gene programs (**Figure 2H, Supplementary Table S2**).

**Figure 2:**
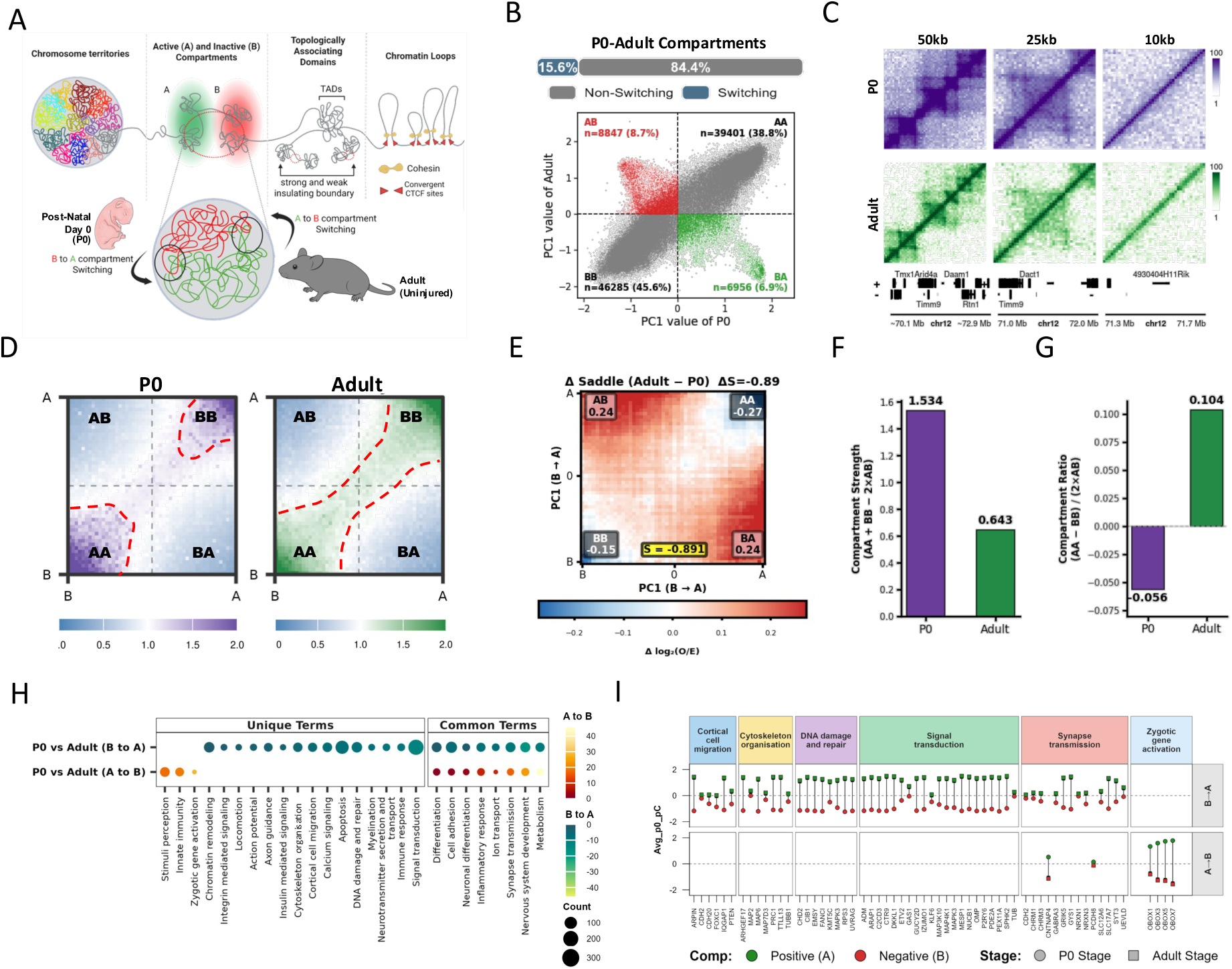
Postnatal maturation reshapes chromatin compartment organization and attenuates global compartment segregation. (A) Hierarchical Organization of the Genome. Schematic representation of 3D genome architecture, illustrating chromosome territories, active (A) and inactive (B) compartments, Topologically Associating Domains (TADs), and chromatin loops stabilized by Cohesin and CTCF. The inset highlights the dynamic switching between A and B compartments.(B) Compartment Switching Dynamics. Scatter plot comparing Principal Component 1 (PC1) values between P0 and Adult stages. Gray points represent stable regions; green points indicate A-to-B compartment switching (8.7%), and red points indicate B-to-A switching (6.9%) during maturation.(C) Multi-resolution Hi-C Contact Maps. Representative heatmaps of chromatin interactions at 50kb, 25kb, and 10kb resolutions for P0 (purple) and Adult (green) stages. Stronger plaid patterns indicate more pronounced compartmentalization.(D) Aggregated Compartment Analysis. saddle plots for P0 (purple) and Adult (green) stages showing the enrichment of contacts within and between A and B compartments. Red dashed lines delineate the boundaries of preferential homotypic (AA and BB) interactions.(E) Differential Saddle Plot. Heatmap representing the change in interaction strength (Δ) between Adult and P0 stages. The scale indicates the gain or loss of specific compartment interactions over time.(F) Quantification of Compartment Strength. Bar plot showing global compartment strength for P0 and Adult stages, calculated as:Strength = (AA + BB)-(2 *AB). A decrease in strength is observed in the Adult stage compared to P0.(G) Compartment Enrichment Ratio. Bar plot illustrating the ratio of homotypic interactions between stages, calculated as: Ratio = (AA - BB) / 2(AB) The shift from negative (P0) to positive (Adult) values indicates a developmental reorganization of A and B densities.(H) Functional Enrichment of Switched Compartments. Dot plot showing Gene Ontology (GO) terms for regions undergoing compartment shifts. Unique and common terms are shown for B-to-A (green) and A-to-B (red) transitions. Dot size represents gene count and color intensity represents statistical significance (-log10 P).(I) Gene-Specific Compartment Shifts. Profile plot of representative genes across functional categories (e.g., Synapse Transmission, Zygotic Activation). Green circles/squares indicate Positive (A) values and red circles/squares indicate Negative (B) values. Circular markers represent P0 stage and square markers represent Adult stage.

Together, these findings demonstrate that postnatal cortical maturation involves coordinated spatial reorganization that consolidates mature neuronal identity while progressively restricting growth-associated chromatin environments.

### Injured cortical neurons show substantial reorganization of three-dimensional genome architecture

At seven days post-injury, transcriptional profiling reveals minimal activation of canonical regeneration-associated genes, yet chromatin accessibility analyses show increased accessibility at select pro-growth regulatory regions, suggesting epigenetic priming that falls short of full transcriptional activation^25,30^. Where three-dimensional genome organization fits within this spectrum remained unknown.

Comparison of PC1 values between uninjured adult and injured cortex revealed that 5.7% of the genome underwent compartment switching following injury, with 3.1% (n=3,116 bins) shifting from B to A and 2.6% (n=2,677 bins) from A to B, against stable fractions of BB (n=50,561, 49.8%) and AA (n=45,140, 44.5%) (**Figure 3B, Supplementary Table S3**). Injury-induced switching thus represents approximately 36.5% of the magnitude observed during developmental maturation, a substantial reorganization triggered solely by a thoracic lesion distal to the cortex. Saddle plot analysis confirmed a marked increase in homotypic interactions and depletion of A-B contacts (**Figure 3D**), with global compartment strength increasing from 0.643 in the adult to 2.167 following injury and delta-S of +1.52 (**Figure 3E,F**). The compartment enrichment ratio decreased from 0.104 to 0.041, reflecting increased B-compartment homotypic interaction density (**Figure 3G**), with this strengthening driven predominantly by intra-chromosomal interactions rather than wholesale repositioning between chromosomes (**Supplementary Figure S3C,D**).

**Figure 3:**
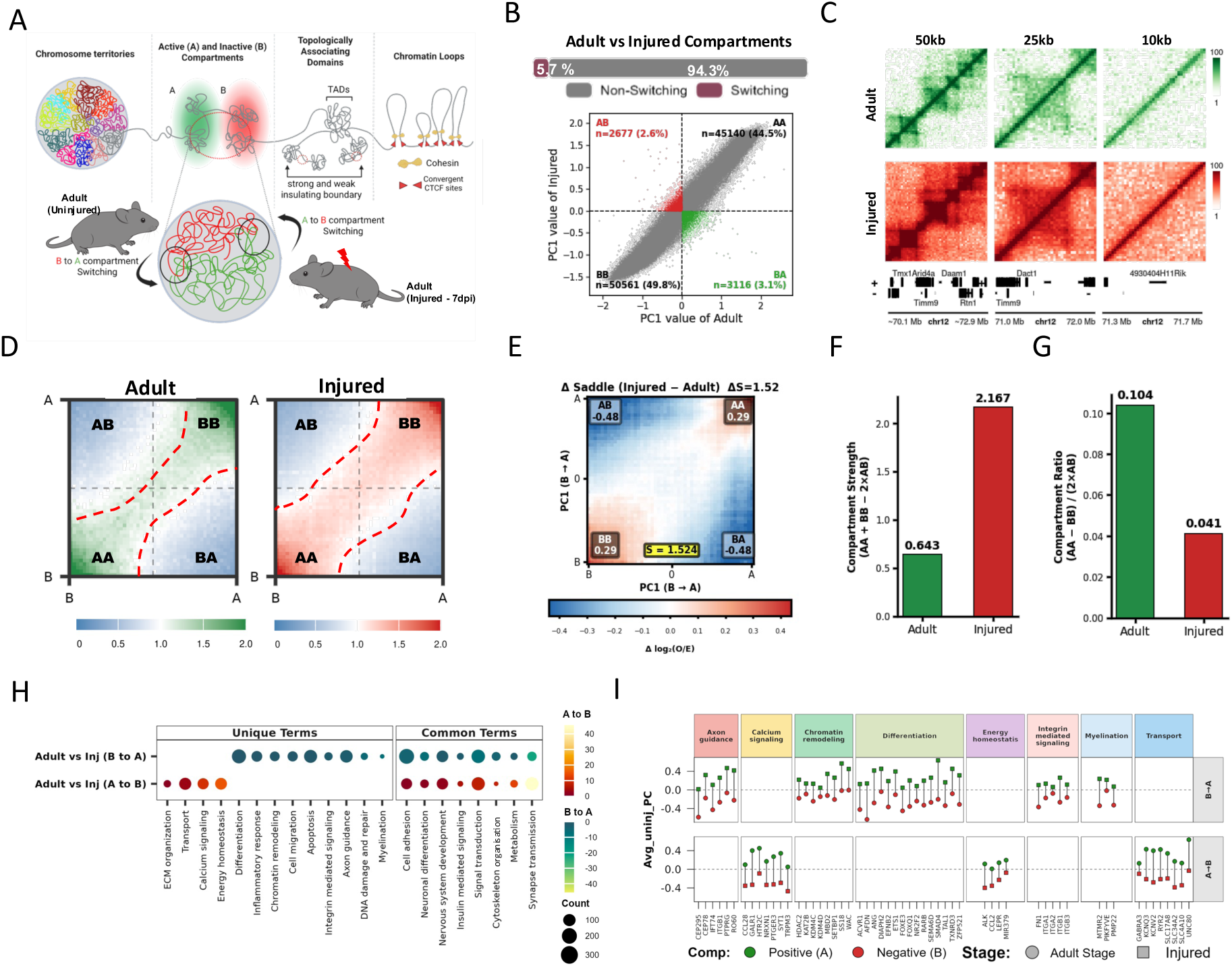
Injured cortical neurons show substantial reorganization of three-dimensional genome architecture. (A) Hierarchical Organization of the Genome. Schematic representation of 3D genome architecture, illustrating chromosome territories, active (A) and inactive (B) compartments, (B) Injury-Associated Compartment Switching. Scatter plot showing the correlation of PC1 values between Adult (Uninjured) and Injured conditions. Gray points indicate stable genomic regions; red points represent regions switching from A-to-B (2.6%), and green points indicate regions switching from B-to-A (3.1%) following injury. (C) Hi-C Contact Map Visualization. Representative Hi-C interaction heatmaps at 50kb, 25kb, and 10kb resolutions for Adult (green) and Injured (red) states. The “plaid” pattern intensity illustrates the degree of spatial segregation between A and B compartments. (D) Aggregated Saddle Plot Analysis. Saddle plots for Adult and Injured states showing the strength of homotypic (AA, BB) versus heterotypic (AB) interactions. (E) Differential Compartmentalization. Heatmap showing the ΔS (Injured - Adult), representing the net change in interaction frequencies between compartments. The scale indicates the gain or loss of specific compartment interactions over time.(F) Global Compartment Strength. Bar plot comparing the overall compartment strength between Adult (0.643) and Injured (2.167) conditions, calculated using the formula: (AA + BB) - (2*AB). (G) Compartment Ratio Comparison. Bar plot showing the ratio of compartment enrichment = (AA + BB)/(2 *AB). (H) Dot plot of Gene Ontology (GO) terms associated with switched compartments. Unique and common terms are shown for B-to-A (red) and A-to-B (green) transitions. Dot size indicates gene count, and color scale represents the significance of enrichment. (I) Gene-Level Compartment Transitions. Profile plot showing PC1 values for specific genes involved in regeneration-related. Green markers denote the active A compartment (positive PC1) and red markers denote the inactive B compartment (negative PC1). Circular markers represent the Adult stage, while squares represent the Injured stage.

Regions gaining active compartment identity were enriched for axon guidance, chromatin remodeling, DNA damage response, differentiation, and myelination pathways, while regions shifting toward repression were enriched for calcium signaling and metabolic processes (**Figure 3H, Supplementary Table S3**). At the gene level, pro-growth regulators including *Epha8*, *Rgs3*, *Coro2b*, and *Cacna1g* shifted toward the active compartment, while genes supporting mature neuronal identity including *Gpm6a*, *Nfia*, and *Zbtb33* transitioned toward repression (**Figure 3I, Supplementary Table S3**). To further resolve the functional hierarchy of these transitions, compartment switches were stratified by the magnitude of PC1 value difference into strong (>1), moderate (0.5–1), and weak (0–0.5) transitions for both the P0-to-adult and adult-to-injured comparisons (**Supplementary Figure S4**). Upon injury, strong B-to-A transitions were enriched for lipid metabolic processes, moderate transitions for cell-cell adhesion, and weak transitions for cell differentiation, while strong A-to-B transitions captured neurogenesis repression. During maturation, strong switching activated cell differentiation and chromatin organisation programs, with inactivated regions enriched for signal transduction and nervous system development, revealing that the functional identity of compartment switches is shaped by both the direction and magnitude of chromatin reorganisation across developmental and injury contexts.

Together, these findings demonstrate that spinal cord injury induces non-trivial three-dimensional genome reorganization in post-mitotic cortical neurons, representing a structural priming mechanism that precedes full transcriptional regeneration.

### Postnatal neuronal maturation builds insulated topological domains that are selectively weakened upon injury

TADs were identified using the HOMER pipeline at 3 kb resolution with 15 kb window, with boundaries defined by local minima in insulation scores. Domains were classified as conserved, shifted (subclassified as strong or weak based on insulation profiles), or unique to each condition (**Figure 4A,D, Supplementary Table S4**). Comparison between P0 and adult cortex revealed 2,138 conserved, 934 shifted, 483 P0-unique, and 1,475 adult-unique domains (Figure 4B, Supplementary Table S4). Postnatal maturation increased total TAD number by 27.9% and reduced median domain size from 369 kb to 171 kb (-53.7%; p<2.1×10⁻¹⁶⁶; Supplementary Figure S5A-C), reflecting fragmentation of broad neonatal domains into smaller insulated neighborhoods. Boundary strengthening predominated during maturation, with 718 TADs gaining insulation compared to 193 weakening (**Figure 4E, Supplementary Table S4**), and genes within strengthened domains were enriched for synaptic transmission, ion transport, and calcium homeostasis pathways (**Figure 4F, Supplementary Table S4**).

**Figure 4:**
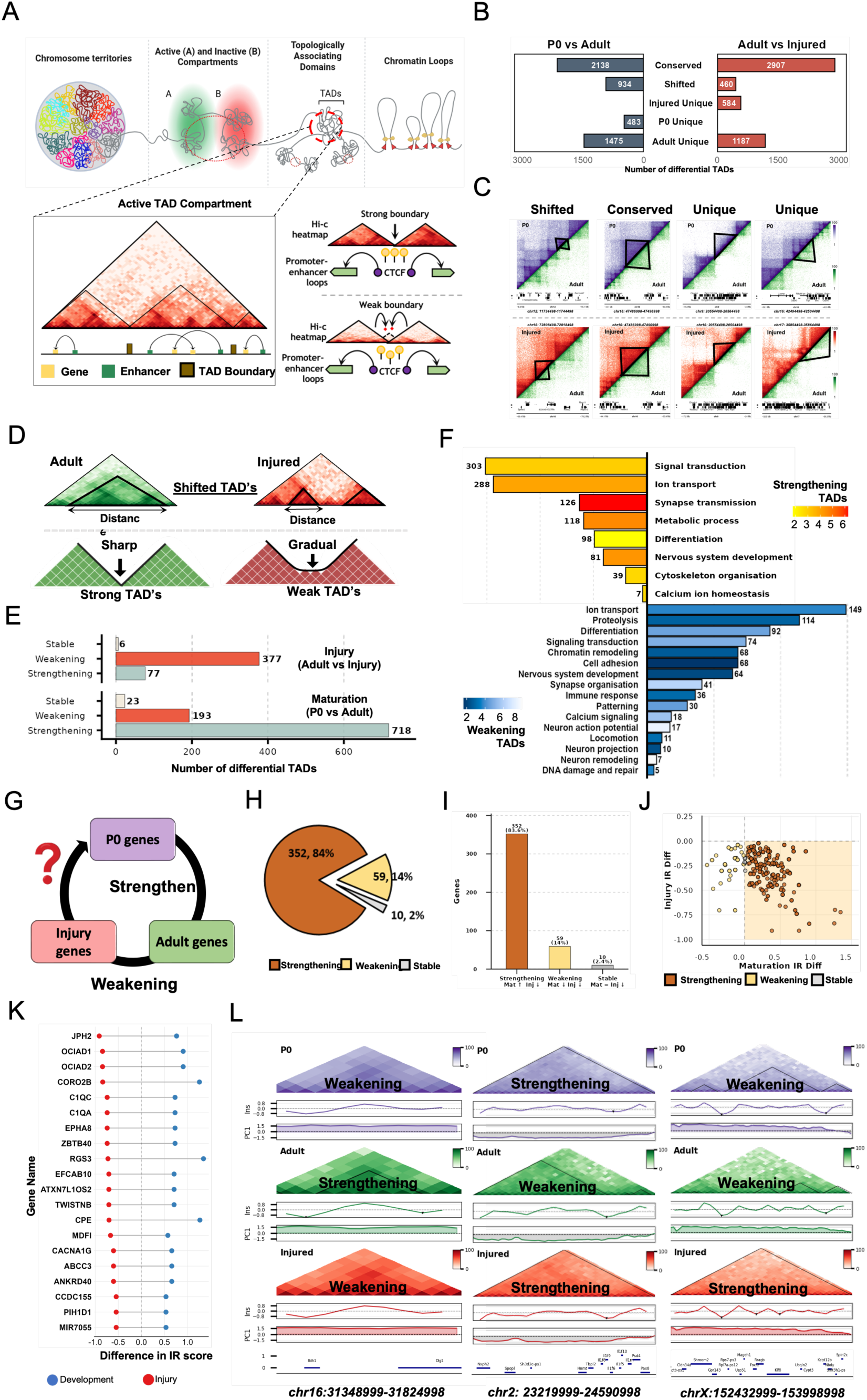
Postnatal neuronal maturation builds insulated topological domains that are selectively weakened upon injury. (A) Schematic illustrating hierarchical chromatin organization, from chromosome territories and A/B compartments to Topologically Associating Domains (TADs) and chromatin loops. Strong TAD boundaries correlate with robust CTCF occupancy and well-defined promoter-enhancer loops, while weak boundaries show diffuse contact signals and reduced insulation. (B) Bar charts showing TADs classified as conserved, shifted, or condition-unique in pairwise comparisons between P0 vs. adult (left) and adult vs. injured (right).(C) Representative Hi-C contact maps illustrating shifted, conserved, and unique TADs across developmental and injury comparisons.(D) Schematic illustrating classification of shifted TADs based on insulation ratio (IR) strength as strong or weak.(E) Bar charts showing shifted TADs classified as stable, weakening, or strengthening in injury (adult vs. injured) and development (P0 vs. adult).(F) Gene Ontology (GO) biological process enrichment for genes within weakening (blue) and strengthening (yellow-red) TADs. Bar length reflects gene count; color intensity indicates – log10 enrichment score. Only terms with p < 0.05 (Bonferroni-corrected) are shown.(G) Schematic posing the central question: whether TADs that strengthen during development (P0 to adult) overlap with TADs that weaken following injury.(H–I) Pie chart and bar chart demonstrating that the majority of genes within TADs weakening after injury (84%, n=352) reside in TADs that had strengthened during postnatal development, with a smaller subset weakening in both contexts (14%, n=59) or remaining stable (2%, n=10).(J) Scatter plot comparing insulation ratio differences of shifted TADs between developmental (x-axis) and injury (y-axis) contexts, colored by classification.(K) Lollipop plot showing IR score differences for the top 20 genes within TADs that strengthen developmentally and weaken after injury.(L) Representative Hi-C contact maps and insulation score profiles across P0, adult, and injured conditions, illustrating TADs that strengthen postnatally and re-weaken upon injury, alongside PC1 and gene tracks.

Following injury, the architectural trajectory reversed. Of all domains, 2,907 were conserved, 460 shifted, and 584 were injured-unique (**Figure 4B, Supplementary Table S4**). TAD number decreased by 13.2% and median domain size expanded from 171 kb to 237 kb (+38.6%; p<7.1×10⁻⁴⁸; **Supplementary Figure S5A-C**), indicating domain coalescence. Boundary weakening predominated, with 377 domains losing insulation compared to only 77 strengthening (**Figure 4E**), and genes within weakening TADs were enriched for chromatin remodeling, nervous system development, neuron remodeling, and DNA damage repair pathways (**Figure 4F, Supplementary Table S4**). Critically, intersection analysis revealed that 84% of genes within TADs weakened by injury resided in TADs that had strengthened during maturation (352 of 421 genes), and scatter plot comparison confirmed a clear anti-correlation between developmental boundary strengthening and injury-induced weakening (**Figure 4G-J**). Lollipop plots of the top 20 genes illustrated this reciprocal directionality at single-gene resolution, including *Jph2*, *Coro2b*, *Rgs3*, *Epha8*, and *Cacna1g* (**Figure 4K**), and representative loci at chr16:31348999-31824998, chr2:23219999-24590998, and chrX:152432999-153998998 confirmed locus-specific reciprocal remodeling (**Figure 4L**).

Together, these findings demonstrate that injury preferentially dismantles the topological architecture consolidated during postnatal development, revealing a non-stochastic directed reorganization that may re-open enhancer-promoter communication at growth-associated loci.

### Injury re-engages neonatal gene programs within reorganized topological domains

We next asked whether injury reconstructs neonatal domain architecture or re-engages similar gene programs through newly configured topological structures(**Figure 5A,Supplementary Table S4**). To determine the functional identity of genes residing within these state-specific domains, we performed gene ontology enrichment analysis on genes located within unique TADs for each condition (**Figure 5B**). P0-unique domains were enriched for neuronal differentiation, axonogenesis, and synapse assembly pathways, consistent with an actively wiring neonatal cortex. Adult-unique domains were enriched for synaptic transmission, ion transport, and neuronal signaling processes, reflecting stabilization of mature cortical circuitry. Injured-unique domains were enriched for axon ensheathment, extracellular matrix adhesion, immune response, and DNA damage signaling pathways, suggesting that injury-associated domains capture transcriptional programs linked to structural remodeling and regenerative signaling.

**Figure 5.**
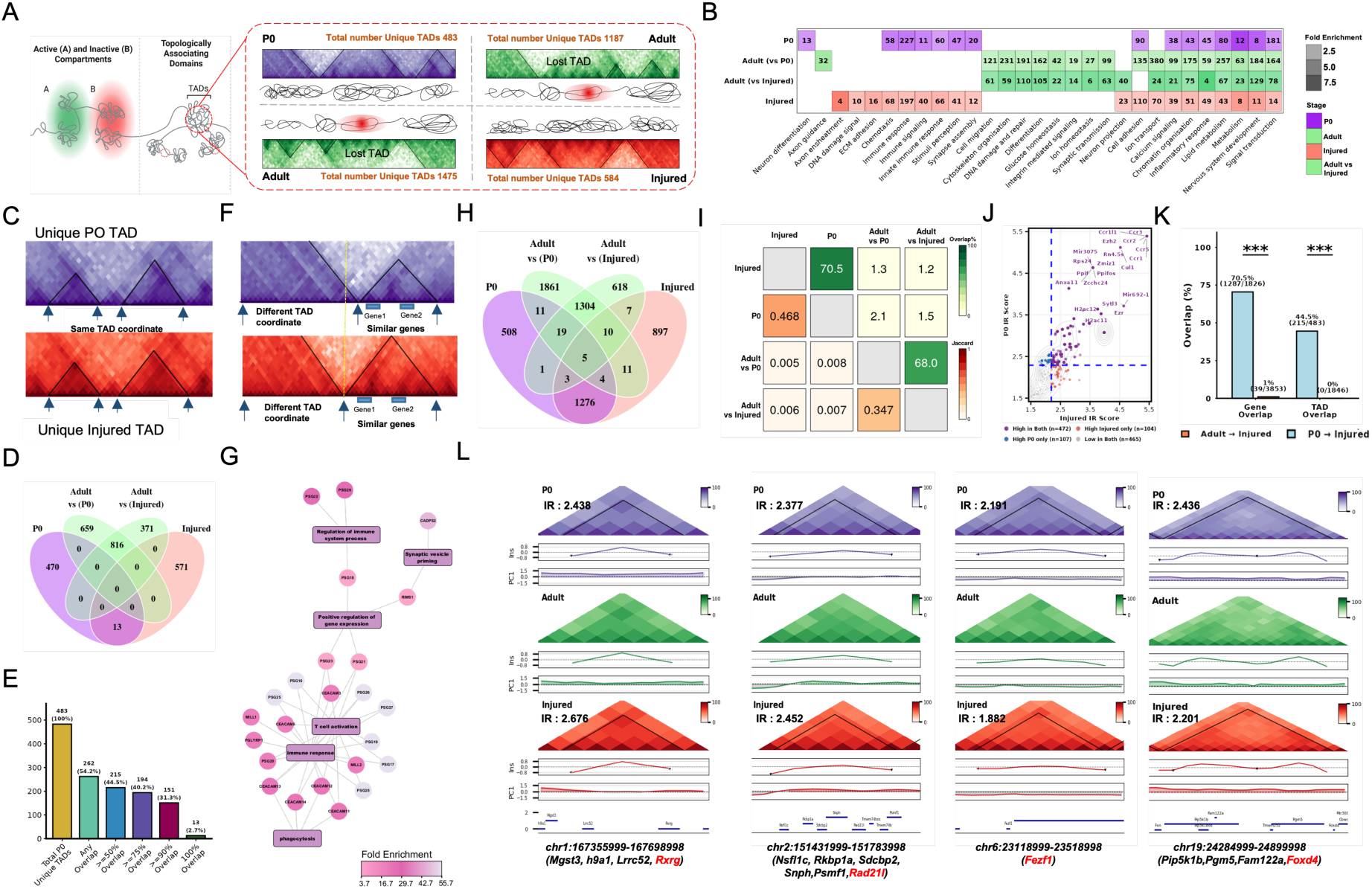
Injury re-engages neonatal gene programs within reorganized topological domains. (A) Schematic illustrating condition-unique TADs in P0 (n=483), Adult (n=1,475), and Injured (n=584) samples, representing domains absent in subsequent datasets.(B) Heatmap displaying GO biological process fold enrichment for genes within condition-specific TADs across P0 (purple), Adult (green), and Injured (salmon) samples. Darker shading indicates higher enrichment.(C) Representative Hi-C contact maps showing spatial overlap of unique TAD coordinates between P0 and Injured conditions. Black triangles mark TAD boundaries.(D) Venn diagram illustrating overlap of TAD coordinates across all samples, highlighting shared and condition-specific domains.(E) Bar graph showing the percentage of P0 TADs retaining positional overlap with Injured TADs at increasing stringency thresholds (any overlap to 100% reciprocal), demonstrating robust spatial conservation between neonatal and injury states.(F) GO network analysis of genes within the 13 TADs uniquely shared between P0 and Injured conditions, with major regulatory hubs including immune response and T cell activation.(G) Contact matrix highlighting that P0 and Injured samples share similar gene content despite differing TAD boundary coordinates, suggesting functional convergence through distinct structural configurations.(H) Four-way Venn diagram illustrating gene overlap within TADs across P0, Adult, and Injured datasets, revealing preferential gene sharing between neonatal and injury states.(I) Jaccard similarity heatmap quantifying gene overlap between samples, with higher values indicating greater genomic similarity. P0 and Injured conditions show the highest similarity relative to Adult.(J) Scatter plot classifying 1,276 genes shared between P0 and Injured based on IR score dynamics: high in both (purple, n=472), high in Injured only (salmon, n=104), high in P0 only (blue, n=107), and low in both (gray, n=465).(K) Bar graph comparing gene and TAD overlap between Adult↔Injured and P0↔Injured, demonstrating significantly greater overlap between P0 and Injured than between Adult and Injured (Fisher’s exact test, ***p<0.001).(L) Representative genomic tracks and contact maps illustrating TAD structure and associated gene content across conditions.

We next asked whether injury recreates neonatal domain architecture at the level of genomic coordinates (**Figure 5C**). Four-way comparison of TAD boundary coordinates revealed that only 13 domains shared identical coordinates between P0 and injured, representing 2.7% of P0-unique and 2.2% of injured-unique TADs, confirming minimal coordinate-level recapitulation (**Figure 5D**).

Gene ontology analysis of these 13 precisely shared domains revealed enrichment for immune response and T cell activation pathways (**Figure 5G, Supplementary Table S4**). Spatial overlap analysis showed that while exact coordinate matching was rare, substantial partial co-occupancy exists between P0 and injured TADs well above chance (**Figure 5E**). Shifting to gene-based comparison revealed a markedly different pattern: 1,276 genes were shared specifically between P0 and injured unique TADs, representing the largest inter-condition overlap in the entire analysis, while adult-to-injured overlap comprised only 7 genes (**Figure 5H**). The Jaccard similarity between P0 and injured was 0.468, substantially higher than adult-to-injured (0.347) or any other cross-condition pair (Figure 5I), and both gene and TAD coordinate overlaps between P0 and injured exceeded adult-to-injured comparisons by Fisher’s exact test (p<0.001) (**Figure 5K**).

Among the 1,276 shared genes having 1148 unique genes, 472 showed high insulation in both P0 and injured states, 104 in injured, 107 in P0, and 465 low in both (**Figure 5J**). Representative loci confirmed direct domain reactivation: at chr1:167355999-167698998 containing *Mgst3* and *Rxrg*, TAD insulation was strong at P0 (IR=2.438), dissolved in the adult, and re-emerged following injury (IR=2.676), with similar patterns at chr2:151431999-151783998 (P0 IR=2.377, Injured IR=2.452), chr6:23118999-23518998 (P0 IR=2.191, Injured IR=1.882), and chr19:24284999-24899998 (P0 IR=2.436, Injured IR=2.201) (**Figure 5L**).

These findings indicate that injury re-engages neonatal gene programs within reorganized topological domains, with coordinate repositioning masking a deeper functional convergence revealed only through gene-level analysis.

### Chromatin loop remodeling across development and injury converges on shared gene targets through repositioned anchors

Differential loop analysis revealed extensive remodeling in both contexts (**Figure 6B**). During development, 589 loops were P0-unique, 351 adult-unique, 48 shifted, and 586 conserved. Following injury, 1,363 loops were injured-unique, only 98 adult-unique, 18 shifted, and 822 conserved, with injured-unique loops outnumbering adult-unique by approximately 14:1. Only 12 loops shared identical coordinates between P0 and injured (0.5%), yet the genes anchored within these loops showed consistently high enrichment scores across both conditions, including *Capn13* (P0=22.8, Injured=31.6), *Abcg2* (P0=19.8, Injured=31.3), *Rgs2* (P0=18.6, Injured=30), and *Rgs6* (P0=18.3, Injured=28.4) (**Figure 6C, Supplementary Table S5**). Gene-based overlap was far more pronounced: Jaccard index for gene overlap between P0 and injured was 0.19 versus near-zero for injured-adult and P0-adult comparisons, with 255 genes shared between P0 and injured loop anchors and minimal adult contribution (**Figure 6D,E, Supplementary Table S5**).

**Figure 6.**
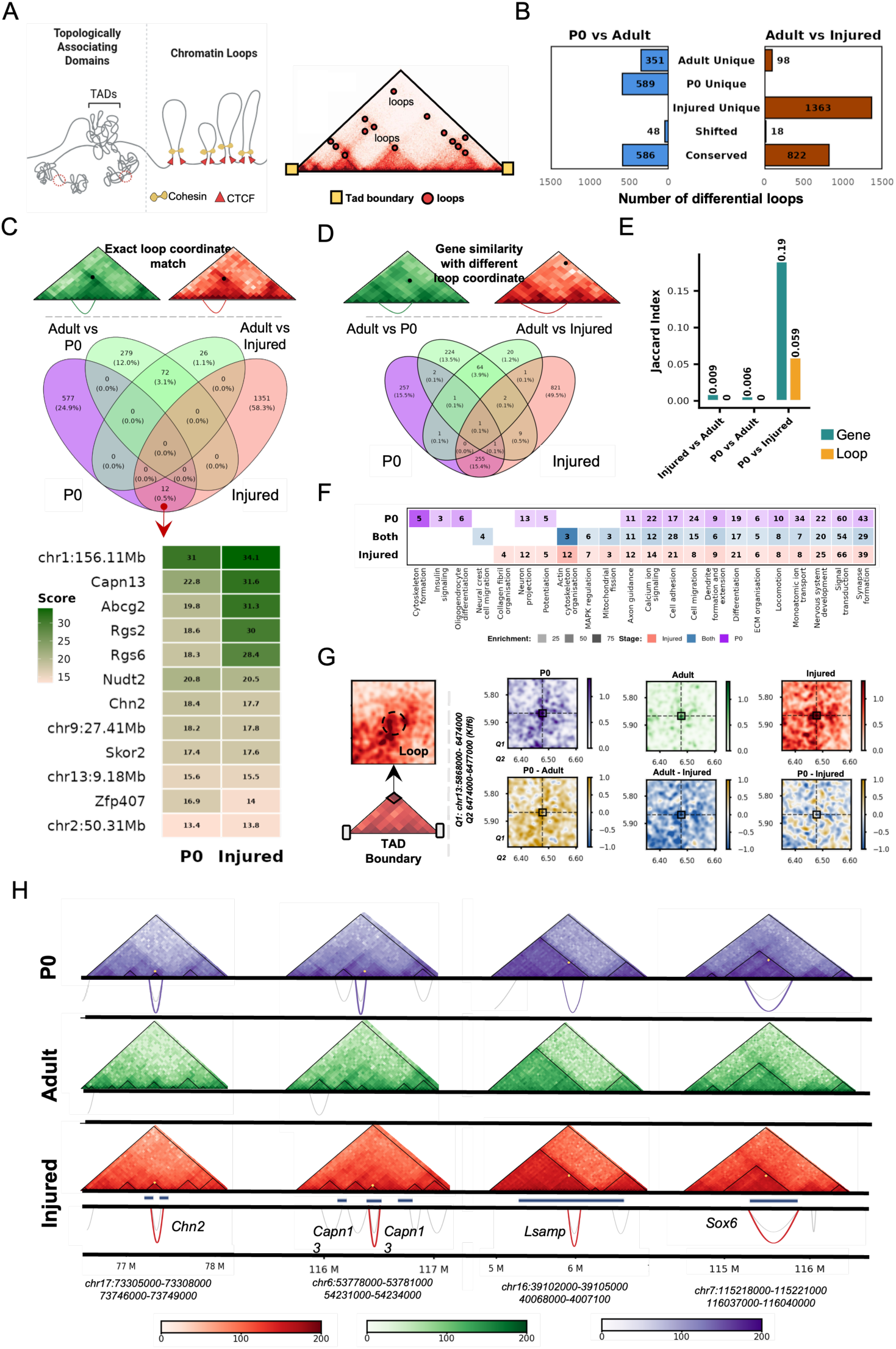
Chromatin loop remodeling across development and injury converges on shared gene targets through repositioned anchors. (A) Schematic of Higher-Order Chromatin Organization. Illustration of Topologically Associating Domains (TADs) and chromatin loops mediated by Cohesin and CTCF. The corresponding Hi-C contact map heatmap shows the representation of TAD boundaries (yellow squares) and loop anchors (red circles). (B) Quantification of Differential Looping. Bar plots showing the number of unique, shifted, and conserved loops (FDR < 0.05) when comparing P0 vs. Adult (blue) and Adult vs. Injured (brown) states. (C) Conservation of Loop Locations. (Top) Schematic of identical loop positions across different contact maps. (Middle) Venn diagram illustrating the overlap of specific chromatin loops across P0, Adult, and Injured datasets at 10 kb resolution. (Bottom) Heatmap showing enrichment scores for specific genes associated with matching unique loop coordinates in P0 and Injured states. (D) Gene-Centric Loop Dynamics. (Top) Schematic showing a gene similarity with different loop coordinate between P0 and injured. (Bottom) Venn diagram showing the overlap of genes associated with chromatin loops, highlighting that while specific loop coordinates may shift, the targeted genes often remain consistent across states. (E) Comparative Similarity via Jaccard Index. Jaccard Index analysis comparing the similarity of genes (cyan) and specific loop coordinates (yellow) between Injured vs. Adult, P0 vs. Adult, and P0 vs. Injured pairs. (F) Functional Enrichment Analysis. Gene Ontology (GO) analysis of genes associated with differential loops (p < 0.05) in P0 (purple), Injured (light orange), and those shared genes between P0 and Injured (blue). (G) TAD Boundary Dynamics with APA score of the *Klf6* Locus. (Left) Illustration of a TAD boundary and loop formation. (Right) Aggregate Peak Analysis (APA) plots centered on the *Klf6* gene (chr13:5.8–6.4 Mb) for P0, Adult, and Injured states. Differential APA plots (bottom row) highlight the gain or loss of looping intensity between stages (P0-Adult, Adult-Injured, and P0-Injured), where darker shading indicates higher loop presence. (H) Genomic Tracks of Shared and Differential Looping. Representative Hi-C contact maps (10 kb resolution) for the *Chn2*, *Capn13*, *Lsamp*, and *Sox6* loci. Tracks highlight conserved loop positions in P0 and Injured states that are absent in the Adult stage (*Chn2*, *Capn13*), as well as genes with shifted anchor points across developmental stages (*Sox6*, *Lsamp*).

Shared genes were enriched for neural crest migration, cytoskeletal organization, axon guidance, and nervous system development (**Figure 6F**). The KLF6 locus illustrated reactivation dynamics directly: looping was present at P0, dissolved in the adult, and re-emerged following injury (**Figure 6G**), positioning KLF6, a transcription factor previously shown to promote corticospinal regeneration^11,12,14,54^, as a candidate for injury-induced reactivation through loop re-engagement. Representative loci at *Chn2* (chr17:73305000-73308000) and *Capn13* (chr6:53778000-53781000) illustrated direct loop re-emergence, while *Lsamp* and *Sox6* (chr7:115218000-115221000) illustrated repositioned anchors converging on the same targets, confirming two modes of injury-induced loop reactivation (**Figure 6H**).

Across all three scales, injury consistently re-engages neonatal gene programs through a reorganized architectural framework, suggesting the motor cortex retains a latent memory of its neonatal regulatory logic that injury partially unlocks.

### Pro-growth gene networks undergo coordinated chromatin reorganization across compartments, domains, and loops

We curated a set of pro-growth genes^13,14,25,55^ and examined their chromatin organization across compartments, TADs, and loops (**Figure 7A, Supplementary Table S6**). Functional network analysis confirmed participation in axon guidance, cytoskeletal organization, neuronal differentiation, chromatin regulation, and regeneration-relevant signaling organized into 12 broad functional clusters (**Figure 7B**). At the compartment level, 16.6% of pro-growth loci underwent switching during development, with stable fractions of AA (n=836, 56.9%) and BB (n=388, 26.4%) and switching fractions of AB (n=109, 7.4%) and BA (n=135, 9.2%) (**Figure 7C**). Following injury, switching was less prevalent at 5.7%, with AA (n=925, 63.0%), BB (n=460, 31.3%), AB (n=36, 2.5%), and BA (n=47, 3.2%) (**Figure 7D**). Of 29 genes undergoing compartment transitions in both contexts, 15 followed a repressive trajectory and 14 followed a reactivation trajectory, with injury selectively restoring active compartment identity at loci repressed during maturation (**Figure 7E**). Representative reactivated genes included *Akap9*, *Kif5c*, *Ptprk*, *Arhgef26*, *Pmp22*, and *Sox4*, while repressed genes included *Fgf13*, *Gpm6a*, *Nfia*, and *Zbtb33* (**Figure 7F,G**).

**Figure 7.**
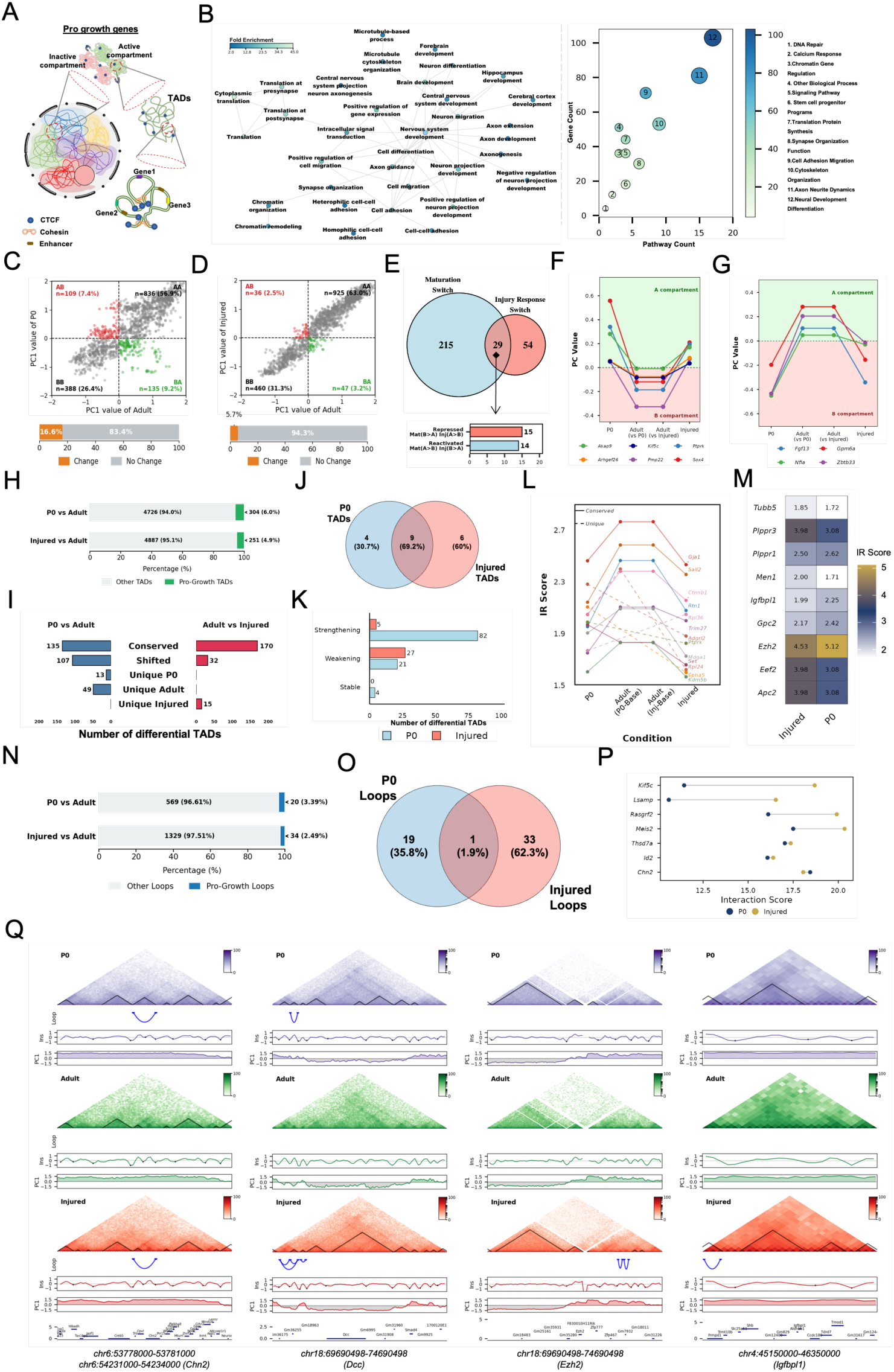
Pro-growth gene networks undergo coordinated chromatin reorganization across compartments, domains, and loops. (A) Schematic illustrating compartment, TAD, and loop organization of pro-growth genes.(B) GO biological process network showing enriched terms associated with pro-growth genes (node size = gene count; edges = shared membership), with a bubble plot summarizing 12 functional pathway categories.(C–D) Scatter plots comparing PC1 values for pro-growth gene regions between P0 vs. adult (C) and adult vs. injured (D), with quadrants indicating compartment transitions (AA, BB, AB, BA).(E) Venn diagram showing overlap between genes undergoing compartment switching during development (n=215) and injury (n=54), with 29 shared genes. Bar chart below quantifies directionality: 15 genes are repressed (B→A developmentally, A→B upon injury) and 14 are reactivated (A→B developmentally, B→A upon injury).(F–G) Line plots showing PC1 trajectories across conditions for representative pro-growth genes that transition from active to inactive compartments during development and return to active upon injury (F; e.g., *Akap9*, *Kif5c*, *Sox4*), and those showing the inverse pattern (G; e.g., *Fgf13*, *Nfia*).(H) Bar chart showing pro-growth TADs as a proportion of all shifted TADs in development (6.0%, n=304) and injury (4.9%, n=251).(I) Bi-directional bar chart classifying pro-growth TADs as conserved, shifted, or condition-unique in developmental and injury comparisons.(J) Venn diagram showing 69.2% (9/13) of P0 pro-growth TADs are shared with injured, indicating strong neonatal-injury structural overlap.(K–M) Bar chart, line plot, and heatmap quantifying and illustrating IR score dynamics of pro-growth TADs across conditions, distinguishing strengthening, weakening, and stable domains.(N) Bar chart showing pro-growth loops as a proportion of all loops in development (3.39%, n=20) and injury (2.49%, n=34).(O–P) Venn diagram revealing that P0 and injured pro-growth loops are largely distinct (only 1 shared loop), with a lollipop plot comparing interaction scores at loop anchors for selected genes including *Kif5c*, *Meis2*, and *Chn2*.(Q) Representative Hi-C contact maps (10 kb) for *Chn2*, *Dcc*, *Igfbpl1*, and *Ezh2* loci illustrating shared and differential looping and TAD organization.

Pro-growth TADs accounted for 6.0% of all TADs in the P0-adult comparison (n=304/4,726) and 4.9% in the adult-injured comparison (n=251/4,887), disproportionate to their genomic fraction (**Figure 7H**). Of pro-growth TADs identified in the neonatal cortex, 69.2% were also present in the injured cortex, with 9 of 28 shared, 4 unique to P0, and 6 unique to injured (**Figure 7I,J**). Directional asymmetry mirrored genome-wide patterns: during maturation 82 pro-growth TADs strengthened and 21 weakened, while following injury 27 weakened and only 5 strengthened (**Figure 7K**). Conserved TADs including *Gja1*, *Sall2*, *Ctnnb1*, and *Rtn1* maintained elevated IR scores across conditions, while unique TADs including *Adgrl2*, *Ptprk*, *Epha5*, and *Kdm5b* showed dissolution in the adult and partial re-emergence following injury, with comparable IR scores between P0-unique and injured-unique shared genes (**Figure 7L,M**). At the loop level, pro-growth loops represented 3.39% of all loops in the P0-adult comparison (n=20/589) and 2.49% in the adult-injured comparison (n=34/1,363), with only 1 loop shared between P0 and injured (1.9%), 19 P0-unique (35.8%), and 33 injured-unique (62.3%) (**Figure 7N,O**). Gene-specific interaction scores revealed that *Kif5c* and *Lsamp* were higher at P0, *Rasgrf2*, *Meis2*, and *Chn2* higher in injured, and *Thsd7a* and *Id2* comparable across states (**Figure 7P**). Representative genomic tracks at *Chn2*, *Dcc*, *Ezh2*, and *Igfbpl1* confirmed gene-specific loop remodeling with P0-injured loop conservation absent in adult (**Figure 7Q**).

Collectively, these findings reveal that adult cortical neurons retain a latent architectural memory of growth that can be structurally re-engaged but remains functionally incomplete, raising the possibility that pro-growth transcription factors could potentiate this latent state beyond the threshold required for regenerative gene activation.

### Successful regeneration engages deeper developmental states of three-dimensional genome architecture

Our previous analyses showed that spinal cord injury partially re-engages developmental chromatin organization without fully restoring architectural features associated with growth competence. Our earlier work revealed that NR2F6 binds a large fraction of growth-associated enhancers^13^, motivating examination of whether it reshapes higher-order genome organization. We therefore asked whether NR2F6, a transcription factor that promotes corticospinal regeneration after severe spinal cord injury, drives a deeper developmental reversion than injury alone, specifically toward the postnatal P0 or more developmentally plastic E12.5 embryonic state (**Figure 8A**).

**Figure 8:**
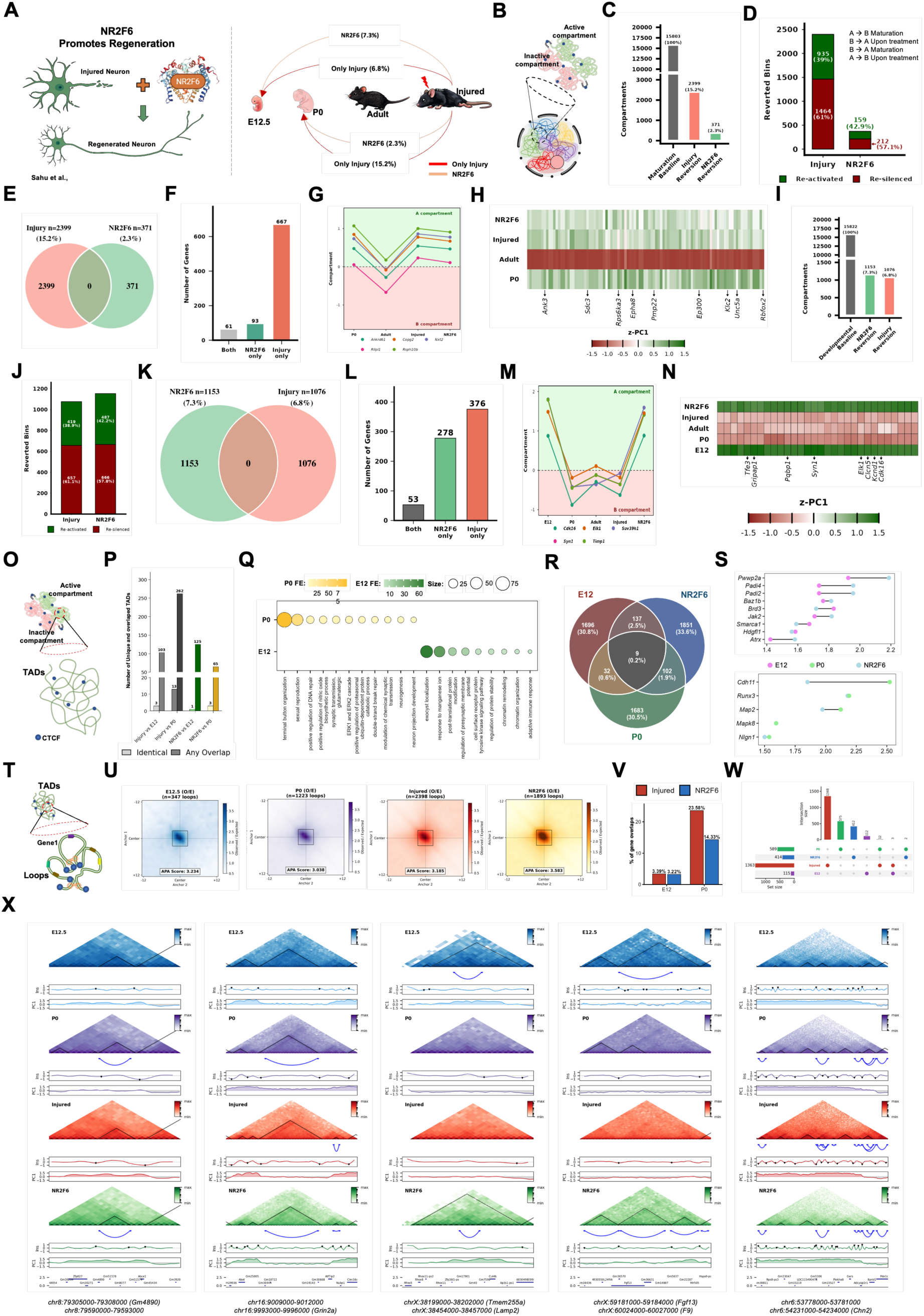
NR2F6 Drives Compartment Reversion Toward Embyonic Chromatin States. (A) Schematic illustrating that injury partially reverts the adult motor cortex chromatin landscape toward a neonatal state, and NR2F6 overexpression further enhances this reversion. Reversion percentages: NR2F6 (7.3%), injury relative to P0 (6.8%), NR2F6 relative to P0 (2.3%), and injury relative to the developmental baseline (15.2%).(B) Schematic of chromatin compartment analysis framework (C–N). (C–E) Relative to the maturation baseline (P0→Adult; 15,803 bins), injury reverts 2,399 bins (15.2%) and NR2F6 reverts 371 bins (2.3%). Both conditions show similar A/B switching proportions (∼42% re-activated, ∼58% re-silenced), with no overlapping reverted bins between injury and NR2F6.(F–H) Bar plot, trajectory plot, and heatmap characterizing genes undergoing compartment switching, distinguishing those shared between NR2F6 and injury from condition-unique genes, and illustrating their compartment dynamics across P0, Adult, Injured, and NR2F6 states.(I–K) Relative to the E12 developmental baseline (1,588 bins), NR2F6 reverts 1,153 bins (7.3%) and injury reverts 1,076 bins (6.8%), again with no bin overlap between conditions, suggesting independent but convergent reversion mechanisms.(L–N) Bar plot, trajectory plot, and heatmap quantifying and visualizing compartment dynamics across the full E12→P0→Adult→Injured/NR2F6 spectrum. NR2F6 uniquely reverts 278 genes, injury uniquely reverts 376 genes, and 53 genes are shared.(O) Schematic of TAD analysis framework. (P–S) Bar plot quantifying TAD conservation between NR2F6/Injured and E12/P0 conditions. GO enrichment dot plot highlights pathways associated with NR2F6-reverted TADs. Venn diagram shows gene overlap within conserved TADs across E12, NR2F6, and P0. Dumbbell plot displays IR scores for selected genes across developmental and treatment conditions.(T) Schematic of chromatin loop analysis framework. (U–W) APA plots reveal strongest loop interactions in NR2F6 (score: 3.583, n=1,893) compared to E12, P0, and Injured. Bar and UpSet plots quantify loop overlap across conditions, highlighting NR2F6-specific and shared loop conservation with neonatal states.(X) Representative Hi-C contact maps (10 kb) for *Gm4890*, *Grin2a*, *Tmem255a*, *Lamp2*, *Fgf13*, *F9*, and *Chn2* loci illustrating shared compartments, TADs, and loops across conditions.

To quantify developmental chromatin reversion, we defined genomic bins switching A/B compartment identity from P0 to adult, establishing a developmental baseline of n=15,803 bins (**Figure 8B,C, Supplementary Tables S2,3,7**). Injury reverted 2,399 bins (15.2%) and NR2F6 reverted 371 bins (2.3%) (**Figure 8C**). Despite injury producing more reverted bins, directionality differed importantly: injury was biased toward re-silencing at 61% re-silenced versus 39% re-activated, while NR2F6 showed a more re-activation-biased profile at 42.9% re-activated versus 57.1% re-silenced (**Figure 8D**), consistent with NR2F6 preferentially restoring active chromatin identity at loci silenced during maturation. Overlap analysis revealed complete non-redundancy at the bin level, with 2,399 injury-unique bins, 371 NR2F6-unique bins, and no shared bins (**Figure 8E**). Gene-level separation was equally complete, with 687 injury-unique genes, 83 NR2F6-unique genes, and only 4 shared, confirming NR2F6 engages an entirely distinct subset of developmentally regulated loci inaccessible to injury signaling (**Figure 8F**). Representative trajectories for *Il1rapl1*, *Atrx*, *Fgf16*, and *Foxo4* confirmed condition-specific compartment dynamics (**Figure 8G,H**).

To extend this analysis to an embryonic reference, we incorporated publicly available Hi-C data from E12.5 cortex generated by the ENCODE Consortium^56^. E12.5 exhibits lower compartmentalisation strength, broader TAD domains, and weaker aggregate loop interactions than P0, consistent with a more plastic chromatin configuration preceding neuronal identity consolidation (**Supplementary Figures S1–S2**). Using E12.5-to-adult transitions as the developmental baseline of n=15,822 bins, injury reverted 1,076 bins (6.8%) while NR2F6 reverted 1,153 bins (7.3%) (**Figure 8I, Supplementary Table S7**). This rank-order reversal, in which NR2F6 transitions from smaller P0 to larger E12 reversion, is the most functionally significant finding of this analysis, indicating NR2F6 preferentially targets embryonic rather than neonatal compartment configurations. Directional analysis again distinguished the conditions: injury remained re-silencing biased at 61.1% versus 38.8% re-activated, while NR2F6 showed 42.2% re-activated versus 57.8% re-silenced (**Figure 8J**). Bin-level overlap remained entirely absent between conditions (**Figure 8K**), and gene-level comparison revealed 376 injury-unique, 278 NR2F6-unique, and 53 shared genes (**Figure 8L**). Representative loci including *Tfe3*, *Clcn5*, *Syn1*, *Elk1*, and *Cdk16* confirmed NR2F6 shifts genes toward developmental compartment identities not reached by injury alone (**Figure 8M,N**).

TAD conservation analysis revealed that NR2F6 exhibited greater structural similarity to E12.5 than to P0 in both the proportion of identical TADs and partial boundary overlap, with insulation ratio profiles showing intermediate scores between E12 and P0 configurations consistent with a hybrid neonatal-embryonic domain architecture (**Figure 8O,P,S**). Gene ontology analysis of conserved TADs revealed enrichment for developmental regulation and neuronal remodeling pathways (**Figure 8Q**), and Venn analysis confirmed NR2F6 contributes 33.6% of conserved TAD genes compared to 30.8% and 30.5% of E12 and P0 (**Figure 8R, Supplementary Table S7**). At the loop level, NR2F6 produced the strongest looping interactions across all conditions with an APA score of 3.583 across n=1,893 loops, exceeding injured (APA=3.185, n=2,398), P0 (APA=3.038, n=1,223), and E12.5 (APA=3.234, n=347) (**Figure 8U**).

NR2F6-associated loops showed greater similarity with developmental states than injured loops alone (**Figure 8V**), and loop intersection analysis confirmed the non-redundant loop repertoire engaged by NR2F6 relative to injury (**Figure 8W**). Representative genomic tracks at *Gm4890*, *Grin2a*, *Tmem255a*, *Lamp2*, *Fgf13*, *F9*, and *Chn2* confirmed that NR2F6 recapitulates developmental contact intensity, domain insulation, and loop engagement at a depth that injury alone cannot achieve (**Figure 8X**).

Together, NR2F6 does not merely intensify the chromatin response triggered by injury but redirects adult cortical chromatin architecture toward earlier developmental configurations across all three scales of genome organization, engaging non-overlapping loci with stronger re-activation bias and superior loop formation. While injury partially re-engages neonatal regulatory architecture, NR2F6 extends this reversion toward a more embryonic-like three-dimensional genome state, suggesting the three-dimensional genome encodes a hierarchy of regenerative potential in which injury primes the first layer of neonatal architectural memory while regeneration-promoting transcription factors access the embryonic chromatin states that fully enable the growth program.

## Discussion

Axon regeneration in the adult mammalian CNS is constrained by an intrinsic failure of injured neurons to re-engage developmental growth programs. Although transcription factor overexpression, epigenetic manipulation, and signaling pathway activation can partially restore regenerative competence, the mechanisms limiting these interventions remain poorly understood. Our findings suggest these limitations arise not only from local chromatin accessibility but from the higher-order organization of the genome. By mapping chromatin compartments, TADs, and loops across developmental and injury states, we show that regenerative potential in corticospinal neurons is encoded within a multi-scale three-dimensional genome architecture that progressively stabilizes during neuronal maturation and can be partially reversed after injury.

Postnatal neuronal maturation is accompanied by extensive compartment reorganization and TAD boundary strengthening, with approximately 15.6% of the genome switching compartment identity as neurons transition from a growth-permissive neonatal to a growth-restricted adult state. These results align with studies showing that neural differentiation and postnatal brain development involve large-scale 3D genome rewiring that proceeds independently of sensory experience, indicating that architectural consolidation is intrinsic to neuronal maturation rather than a passive consequence of reduced activity^51,57,58^, and that post-mitotic neurons continue restructuring their 3D genome across the lifespan^59–61^. Critically, the compartments, TAD boundaries, and loops that consolidate during maturation are specifically enriched at pro-growth gene loci, linking these architectural transitions directly to the loss of regenerative competence rather than reflecting generic genome-wide stabilization. The progressive insulation of regulatory neighbourhoods during maturation is consistent with studies showing that enhancer-promoter contacts become more predictive of transcriptional output during later differentiation^62–64^, effectively locking regulatory architecture into a configuration that restricts reactivation of developmental growth genes.

Spinal cord injury partially reverses these maturation-associated changes, inducing compartment switching and selectively dismantling TAD boundaries strengthened during postnatal development. The magnitude of this response is substantial, with injury-induced compartment reversion representing roughly one-third of the developmental changes that occur during maturation. Given that corticospinal neurons are post-mitotic and physically distant from the injury site, this scale of architectural reorganization is striking and consistent with work showing that post-mitotic neurons can dynamically restructure their 3D genome in response to stimulation, and that neuronal DNA damage associates with multiscale disruption of genome organization^65,66^.

Importantly, injury does not reconstruct the neonatal 3D genome exactly: neonatal loop coordinates are rarely restored, yet the same gene targets are re-engaged through repositioned regulatory anchors, consistent with developmental studies showing that regulatory logic is more conserved than precise geometric configuration^63,67^. Injury therefore approximates developmental regulatory architecture without recreating it, producing a partially permissive chromatin environment that primes but does not fully activate growth-associated gene programs. This dissociation is well supported by the Activity-by-Contact model, in which enhancer output depends multiplicatively on both contact frequency and enhancer activation state^68^: architectural permissiveness is established in the first week after injury, but without sufficient enhancer activation to cross the threshold for productive transcription.

A central implication of these results is that adult cortical neurons retain a latent architectural memory of their developmental growth state. Rather than creating new regulatory architecture, injury exposes a pre-existing developmental regulatory logic embedded within neuronal chromatin topology. This parallels observations in other biological systems where cells retain structural memory of prior regulatory states: memory T cells maintain preconfigured chromatin hubs that enable rapid transcriptional recall without constitutive activation^69^, hippocampal engram neurons establish partial enhancer-promoter contacts during memory encoding that precede full transcriptional activation during recall^70,71^, and induced pluripotent stem cells retain epigenetic memory of their tissue of origin that biases subsequent differentiation^72^. Our injury data fit this model precisely, with architectural priming in the first week after injury representing a structurally meaningful intermediate state reflecting partial re-engagement of a stored developmental regulatory topology rather than a failure of the injury response.

NR2F6 reveals an additional dimension to this architectural model. While injury primarily reverts chromatin compartments toward the neonatal P0 state, NR2F6 drives reversion toward the earlier embryonic E12 configuration, engaging entirely non-overlapping genomic loci with a stronger re-activation bias. This rank-order reversal, in which NR2F6 produces a smaller P0 but larger E12 reversion than injury, identifies depth of developmental access rather than scale of architectural change as the functionally relevant variable for regenerative outcome. This hierarchical pattern resonates with stem cell and immune memory biology, where access to earlier developmental chromatin states confers increased plasticity^73,74^, and with observations that histone H1 loss and hematopoietic development both demonstrate that accessing earlier developmental architecture increases regulatory competence in ways that shallower changes cannot achieve^75,76^. Furthermore, NR2F6 produces the strongest loop formation across all conditions, with an APA score of 3.583 exceeding injured, P0, and E12.5. Given that the Activity-by-Contact model predicts enhancer output is multiplicatively determined by contact frequency and enhancer activation state^68^, NR2F6-induced loop strengthening combined with its known enhancer-binding activity at growth-associated loci may together cross the threshold for productive transcriptional activation that injury-induced partial contact restoration cannot reach alone.

Our findings reframe the chromatin barrier to CNS regeneration as a topological problem^39,77^. Even when local accessibility is restored, the three-dimensional context of a locus, its compartment identity, TAD insulation, and loop connectivity, determines whether enhancers can productively engage their target promoters^37^. This explains why broad chromatin accessibility interventions can produce widespread remodeling without regeneration, while NR2F6, which accesses deeper developmental architectural states, successfully promotes axon growth. Emerging tools for targeted architectural remodeling, including dCas9-based loop induction^78,79^, synthetic CTCF anchor insertion^80^, and compartment-switching via heterochromatin redistribution^81^, could in principle be directed at the specific loci re-engaged by NR2F6 but not by injury. More broadly, our finding that the adult motor cortex retains a latent architectural memory of its developmental growth state suggests that the barrier to regeneration is not the absence of regulatory information but its structural inaccessibility, positioning 3D genome topology as an independent and targetable regulatory layer controlling regenerative competence in the adult CNS.

## Materials and Methods

### Animal Husbandry

All animal experiments were conducted in accordance with protocols approved by the Institutional Animal Ethics Committee (IAEC) of the Centre for Cellular and Molecular Biology (CCMB), Hyderabad, India (51/2023, 79/2024, 67/2025). Mice were housed under a 12-hour light/12-hour dark cycle, with controlled environmental conditions maintained at 23–25 °C and 40–60% relative humidity.

### Animal Surgeries – Thoracic Crush

Adult mice were anesthetized and the dorsal cervical–thoracic region was shaved and disinfected prior to surgery. Animals were subsequently secured in a custom spinal stabilization apparatus to minimize movement during the procedure. A longitudinal midline incision extending from the base of the ears to ∼2 cm caudally along the upper back was made to expose the surgical site. Laminectomy was performed at the T10-T12, which was identified by the spinous process. The cord was then subjected to a standardized crush injury using calibrated blunt forceps applied across the full width of the cord for 10 seconds to ensure consistent pressure. After injury, the vertebral elements were repositioned, the musculature and surrounding tissues were restored to their anatomical orientation, and the skin incision was closed with black nylon thread. Post-operative care included administration of analgesics and antibiotics, manual bladder expression twice daily until recovery of function, and maintenance on a heating pad during the immediate recovery period. Successful injury was verified by the presence of hindlimb paralysis following surgery. All procedures were conducted aseptically within a Class II biosafety cabinet.

### High-throughput Chromosome Conformation Capture (Hi-C)

Hi-C libraries were generated from three experimental groups: postnatal day 0 (P0), uninjured adult, and adult mice at 7 days following spinal cord injury, using protocol adapted from the ENCODE Consortium^82^ with minor modifications as previously described by our group^13,25^. The NR2F6 Hi-C dataset was obtained from a preprint from our group^13^ (PRJNA1217058). For all in-house generated libraries, motor cortices (35-40mg, from 5 animals) were isolated from the mouse brain, the tissues were kept on ice and pulverized in liquid nitrogen with pre-chilled mortar and pestle. The powdered tissue was resuspended in 500 µL of 1× PBS, transferred to a 15 mL conical tube, and the volume was adjusted to 5 mL with 1× PBS. Tissues were crosslinked with 1% formaldehyde (Thermo Fisher, 12755) for 20 min at room temperature with shaking (180–200 rpm). Crosslinking was quenched with 2.5 M glycine (MP Biomedicals, 808822) to a final concentration of 0.125 M for 5 min.

The samples were centrifuged at 2000 × g for 15 min, the supernatant was discarded, and the pellet was washed with 500 µL ice-cold 1× PBS followed by centrifugation at 2500 × g for 10 min. The pellet was subjected to nuclei isolation using ice-cold Hi-C lysis buffer (5 mM CaCl₂, 3 mM MgAc₂, 2 mM EDTA, 0.5 mM EGTA, 10 mM Tris-HCl pH 8.0, 1 mM DTT, 0.1 mM PMSF) supplemented with 1× protease inhibitors (MCE, HY-K0010). Nuclei were purified by density gradient centrifugation using 1 M sucrose buffer (1 M sucrose, 3 mM MgAc₂, 10 mM Tris-HCl pH 8.0). The sucrose solution was added carefully along the tube wall to form a gradient, and samples were centrifuged at 1000 × g for 5 min (deceleration: 3; acceleration: 9). The supernatant was removed and the nuclei pellet was resuspended in 50 µL of 0.5% SDS and incubated at 62°C for 10 min. SDS was quenched by adding 10% Triton X-100 followed by incubation at 37°C for 15 min.

Chromatin was digested overnight at 37°C with 100 U of MboI (NEB, R0147) in a thermomixer at 700 rpm. The enzyme was inactivated at 62°C for 20 min. DNA ends were filled in with 0.4 mM biotin-14-dATP (Thermo Scientific, 19524-016), 10 mM each dCTP, dGTP, and dTTP (Thermo Scientific), and DNA Polymerase I Klenow fragment (NEB, M0210) at 37°C for 90 min (500 rpm). Ligation was performed at room temperature for 4 h (300 rpm) in the presence of T4 DNA ligase buffer, 10% Triton X-100, 20 mg/mL BSA (Sigma, A7906), and T4 DNA ligase (NEB, M0202).

For crosslink reversal, Proteinase K (NEB, P8102) was added to a final concentration of ∼0.7 mg/mL with ∼1% SDS and incubated at 55°C for 30 min. NaCl was then added to a final concentration of 0.5 M and samples were incubated overnight at 65°C. DNA was purified by ethanol precipitation method using 1.6× volumes of ice-cold 100% ethanol and 0.1× volume of M sodium acetate (pH 5.2), followed by incubation at −80°C for 45 min and centrifugation at maximum speed for 20 min at 4°C. Pellets were washed with 70% ethanol, air-dried, and resuspended in 130 µL of 10 mM Tris-HCl (pH 8.0). The purified DNA was sheared to 300–500 bp using a Covaris M220 (Peak Incident Power: 50, Duty Factor: 10, Cycles/Burst: 350, Time: 70 s). The volume was adjusted to 200 µL with dH₂O, and size selection was performed using 0.55× SPRI beads (Beckman Coulter, B23318). Biotin-labeled fragments were captured with Streptavidin T1 beads (Invitrogen, 65601).

To repair fragmented ends and remove biotin from unligated ends, the beads were resuspended in 100ul of the solution containing 1X NEB T4 DNA ligase buffer (NEB, B0202), 5ul of 10mM dNTP mix (NEB, N0447S), 5ul of 10U/ul NEB T4 DNA Polymerase (NEB, M0203), 1ul of 5U/ul Klenow (NEB, M0210). The samples were incubated for 30minutes at room temperature and the beads were reclaimed on a magnetic stand. For the addition of dA-tail, the beads were resuspended in 100ul of the solution containing 90ul of 1X NEB Buffer 2, 5ul of 10mM dATP (Invitrogen 100004916) 5ul of 5U/ul Klenow (exonuclease, NEB, M0212). The samples were incubated for 30minutes at 37°C, separated on the magnetic stand and the supernatant was discarded.

Adapters were ligated using Illumina TruSeq adapters (Illumina, 20040870) by resuspending beads in 50 µL of 1× NEB Quick Ligation Reaction Buffer (NEB, B2200S), 2 µL NEB Quick Ligase (NEB, M2200), and 3 µL indexed adapters, followed by incubation at room temperature for 15 min. Beads were washed and resuspended in 50 µL of 10 mM Tris-HCl (pH 8.0). Libraries were PCR-amplified for 11 to 12 cycles directly from T1 beads using Illumina primers according to the manufacturer’s protocol.

Following amplification, beads were separated magnetically and the supernatant was transferred to a new tube. Final size selection was performed using 0.7× AMPure XP beads (Beckman Coulter, A63880) with a 15 min incubation at room temperature. DNA was eluted in 25–50 µL of 10 mM Tris-HCl (pH 8.0). Libraries were sequenced on an Illumina NovaSeq platform to a depth of ∼700 million base pairs.

### Hi-C Data Processing

#### Quality Control and Read Trimming

Raw paired-end Hi-C sequencing reads were subjected to quality control and adapter trimming using fastp (v0.23.2) ^83^. Reads were filtered using a minimum Phred quality score of 30 (--qualified_quality_phred 30), adapter sequences were automatically detected and removed for paired-end data (--detect_adapter_for_pe), and reads shorter than 20 bp after trimming were discarded (--length_required 20). Quality reports were generated in both HTML and JSON formats for each sample. Processing was performed using 10 threads.

#### Read Alignment to the Reference Genome

Following initial quality control, trimmed reads were further processed to remove ligation junction sequences using the HOMER homerTools utility ^84^. Specifically, reads were trimmed at MboI restriction enzyme recognition sites (GATC) using the parameters ‘-3 GATC -mis 0 - matchStart 20 -min 20’, which trims from the 3’ end at GATC motifs with zero mismatches, beginning the search at position 20, and retaining reads of at least 20 bp.

The trimmed reads from each end (R1 and R2) were aligned independently to the mouse reference genome (mm10; GRCm38) using Bowtie2 (v2.4.5) ^85^ in single-end mode with default parameters, using 20 parallel threads (-p 20). Separate SAM files were generated for each read end per sample. The mm10 genome index was pre-built using bowtie2-build prior to alignment.

#### Tag Directory Construction

Aligned reads from both ends were combined and used to construct HOMER tag directories using the makeTagDirectory command^84^. A coverage saturation parameter of -tbp 1 was applied to limit the number of tags per base pair, reducing PCR duplicate bias. Tag directories serve as the primary data structure for all downstream HOMER-based analyses.

#### A/B Compartment Identification

Chromatin compartment analysis was performed using the HOMER runHiCpca.pl script^84^. Principal component analysis (PCA) was carried out at 25 kb resolution with a 50 kb smoothing window (-res 25000 -window 50000) against the mm10 genome. The first principal component (PC1) values were exported as bedGraph files and used to assign genomic bins to A compartments (active, positive PC1) or B compartments (inactive, negative PC1). Analysis was performed using 20 CPU threads.

#### Hi-C Interaction Matrix and Compaction Statistics

Hi-C interaction matrices and insulation/compaction statistics were generated using the HOMER analyzeHiC command ^84^ at 5 kb resolution with a 15 kb window (-res 5000 -window 15000). The -nomatrix flag was used to suppress full matrix output, and -compactionStats auto was applied to compute insulation ratio (IR) scores genome-wide. These scores were used to assess local chromatin compaction at domain boundaries. Analysis was run using 20 CPU threads.

#### TAD and Loop Identification

Topologically associating domains (TADs) and chromatin loops were identified using the HOMER findTADsAndLoops.pl pipeline^84^. Calling was performed at 3 kb resolution with a 15 kb window (-res 3000 -window 15000) using 10 CPU threads and the mm10 genome as reference. TAD boundaries were defined based on the local minimum insulation score, and loops were called based on significant local enrichment of Hi-C contacts relative to the expected background model.

#### Compartment Strength Analysis and Saddle Plots

Compartment strength was quantified and visualised using saddle plots computed with the cooltools Python package (v0.6.1)^86^. Hi-C contact matrices were stored and accessed in .cool format using the cooler library (v0.9.3)^86^. Genomic bin-level PC1 values derived from HOMER were mapped to cooler bins using bioframe (v0.4.1)^87^. Expected cis interaction profiles were computed using cooltools.expected_cis() over the full genome view. Saddle plots were generated using cooltools.saddle() with 50 quantile bins (n_bins=50) and a quantile range of 0.025–0.975 to exclude extreme outlier bins. Compartment strength was computed as the sum of homotypic interactions (AA + BB) minus heterotypic interactions (AB + BA) in the five corner bins of the saddle matrix.

To identify condition-specific compartment changes, differential saddle plots were generated by computing the log2 ratio of observed-over-expected (O/E) interaction matrices between conditions (ΔSaddle = log2[O/E₁] − log2[O/E₂]). Analyses were restricted to genomic regions associated with pro-growth genes, identified from pre-compiled gene lists. All saddle plots were rendered using matplotlib (v3.7)^88^ and saved at 300 dpi.

#### Aggregate Peak Analysis (APA) for Chromatin Loops

Chromatin loop strength was assessed by Aggregate Peak Analysis (APA) using custom Python scripts based on the cooler library^89^. Hi-C contact matrices at 10 kb resolution were fetched from .mcool files. For each annotated loop anchor pair, the genomic coordinates were converted to bin indices and a submatrix of ±25 bins (250 kb) around each loop centre was extracted. Submatrices with missing chromosomal data or extending beyond chromosome boundaries were excluded. Each submatrix was normalised by its mean contact frequency to account for differences in sequencing depth and local interaction background. Valid submatrices were averaged to produce the APA pileup map for each condition. Differential APA maps were computed by subtracting the average pileup of one condition from another. Individual and differential APA plots were visualised with matplotlib^88^ and saved at 300 dpi.

#### Aggregate Domain Analysis (ADA) for TADs

TAD insulation strength was quantified using Aggregate Domain Analysis (ADA) implemented in custom Python scripts using cooler^89^ and scipy^90^. TADs were filtered to retain domains between 100 kb and 2 Mb in size. For each TAD, a submatrix was extracted from the Hi-C contact map encompassing the TAD body plus flanking regions equal to the full TAD length on each side (flank fraction = 1.0). Raw contact counts were log1p-transformed prior to averaging. All submatrices were rescaled to a uniform 50 × 50 pixel grid using bilinear interpolation (scipy.ndimage.zoom, order=1) to allow aggregation across TADs of varying sizes. The ADA score was computed as the mean intra-TAD contact intensity minus the mean flanking contact intensity (upper triangle values only, excluding the diagonal). Averaged matrices were visualised as triangle heatmaps using matplotlib^88^. Differential ADA plots were generated by subtracting averaged matrices between conditions, and dashed vertical lines were overlaid to delineate TAD boundaries. All plots were saved at 300 dpi.

#### Downstream Statistical Analysis and Visualisation

All downstream statistical analyses, including differential compartment classification, TAD boundary comparisons, loop categorisation, and gene overlap analyses, were performed in R (v4.3)^91^. Data were manipulated using the tidyverse suite of packages and visualised using ggplot2 (v3.4)^92^.

Genome-wide Hi-C contact maps were visualised using HiCExplorer (v3.7)^93^ and the cooler Python library^89^. PC1 compartment tracks, insulation ratio scores, and normalised interaction frequency matrices were visualised in R using the plotgardener package (v1.8)^94^, which enabled precise multi-panel genomic figure assembly. Contact maps were plotted with balanced normalisation applied.

All analyses were conducted using the mouse reference genome assembly mm10 (GRCm38), obtained from the UCSC Genome Browser. Chromosome sizes were retrieved from the corresponding genome index files.

## Resource availability

All genomics datasets generated in this study have been deposited in the NCBI Gene Expression Omnibus (GEO) under accession number PRJNA1217518. A detailed summary of datasets and corresponding sample information is provided in Supplementary Table S8. The metadata detailing the Supplementary Tables used for the results is provided in Supplementary Table S1. The code used for data processing and figure generation is available at https://github.com/mano2991/3D-organisation-in-CNS-neurons.

## Acknowledgements

We thank CSIR-CCMB for providing research facilities and support staff. We acknowledge Mr. S. Prasanth and Mr. N. Sai Ram (Animal House), Dr. Karthik Baradwaj, Ms. Tulasi Nagabandi, Dr. Md. Jafurulla (Genomics), and B. Venkateshwarlu for their technical assistance. We also appreciate Fine BioChemical for the timely chemical supplies. We would also like to thank the funding agencies Council of Scientific and Industrial Research (CSIR), Department of Biotechnology (DBT), Anusandhan National Research Foundation (ANRF), and BFI Biome, Government of India.

## Author contributions

Conceptualization and experimental design: A.S.M. M.K.K and I.V. Data curation: A.S.M., M.K.K, F.F., and I.V. Formal analysis: A.S.M., M.K., F.F., and I.V. Investigation: A.S.M. and I.V. Methodology: A.S.M., M.K., F.F., D.S.B., D.K., Y.S., and I.V. Funding acquisition: I.V. Project administration: A.S.M. M.K.K and I.V. Resources: I.V. Software: A.S.M., M.K., and I.V. Supervision: A.S.M. M.K.K and I.V. Validation: A.S.M. and I.V. Visualization: A.S.M., M.K., F.F., and I.V. Writing – original draft: A.S.M. and I.V. Writing – review and editing: A.S.M., M.K.K, F.F., and I.V. All authors read and approved the final manuscript.

## References

1. Huebner, E. A. & Strittmatter, S. M. Axon Regeneration in the Peripheral and Central Nervous Systems. Results Probl Cell Differ 48, 339–351 (2009).

2. Kai Liu et al. Neuronal Intrinsic Mechanisms of Axon Regeneration. Annual Review of Neuroscience 10.1146/annurev-neuro-061010-113723 (2011) doi:10.1146/annurev-neuro-061010-113723.

3. Sun, F. & He, Z. Neuronal intrinsic barriers for axon regeneration in the adult CNS. Curr Opin Neurobiol 20, 510–518 (2010).

4. Mahar, M. & Cavalli, V. Intrinsic mechanisms of neuronal axon regeneration. Nat Rev Neurosci 19, 323–337 (2018).

5. Fawcett, J. W. The Struggle to Make CNS Axons Regenerate: Why Has It Been so Difficult? Neurochem Res 45, 144–158 (2020).

6. Ilaria Palmisano et al. Epigenomic signatures underpin the axonal regenerative ability of dorsal root ganglia sensory neurons. Nature Neuroscience 10.1038/s41593-019-0490-4 (2019) doi:10.1038/s41593-019-0490-4.

7. Chen, M. S. et al. Nogo-A is a myelin-associated neurite outgrowth inhibitor and an antigen for monoclonal antibody IN-1. Nature 403, 434–439 (2000).

8. Qiu, J., Cai, D. & Filbin, M. T. Glial inhibition of nerve regeneration in the mature mammalian CNS. Glia 29, 166–174 (2000).

9. Mukhopadhyay, G., Doherty, P., Walsh, F. S., Crocker, P. R. & Filbin, M. T. A novel role for myelin-associated glycoprotein as an inhibitor of axonal regeneration. Neuron 13, 757–767 (1994).

10. Filbin, M. T. Myelin-associated inhibitors of axonal regeneration in the adult mammalian CNS. Nat Rev Neurosci 4, 703–713 (2003).

11. Kramer, A. A., Olson, G. M., Chakraborty, A. & Blackmore, M. G. Promotion of corticospinal tract growth by KLF6 requires an injury stimulus and occurs within four weeks of treatment. Experimental Neurology 339, 113644 (2021).

12. Wang, Z. et al. KLF6 and STAT3 co-occupy regulatory DNA and functionally synergize to promote axon growth in CNS neurons. Sci Rep 8, 12565 (2018).

13. Sahu, Y. et al. Nuclear Receptor Transcription Factors promote axon regeneration in the Adult Corticospinal Tract. Preprint at 10.1101/2025.06.19.660319 (2025).

14. Venkatesh, I. et al. Co-occupancy identifies transcription factor co-operation for axon growth. Nat Commun 12, 2555 (2021).

15. Norsworthy, M. W. et al. Sox11 Expression Promotes Regeneration of Some Retinal Ganglion Cell Types but Kills Others. Neuron 94, 1112–1120.e4 (2017).

16. Wang, Z., Reynolds, A., Kirry, A., Nienhaus, C. & Blackmore, M. G. Overexpression of Sox11 promotes corticospinal tract regeneration after spinal injury while interfering with functional recovery. J Neurosci 35, 3139–3145 (2015).

17. Dhara, S. P. et al. Cellular reprogramming for successful CNS axon regeneration is driven by a temporally changing cast of transcription factors. Sci Rep 9, 14198 (2019).

18. Venkatesh, I., Mehra, V., Wang, Z., Califf, B. & Blackmore, M. G. D evelopmental C hromatin R estriction of P ro- G rowth G ene N etworks A cts as an E pigenetic B arrier to A xon R egeneration in C ortical N eurons. Developmental Neurobiology 78, 960–977 (2018).

19. Gaub, P. et al. HDAC inhibition promotes neuronal outgrowth and counteracts growth cone collapse through CBP/p300 and P/CAF-dependent p53 acetylation. Cell Death Differ 17, 1392–1408 (2010).

20. Gaub, P. et al. The histone acetyltransferase p300 promotes intrinsic axonal regeneration. Brain 134, 2134–2148 (2011).

21. Franziska Müller et al. CBP/p300 activation promotes axon growth, sprouting, and synaptic plasticity in chronic experimental spinal cord injury with severe disability. PLOS Biology 10.1371/journal.pbio.3001310 (2022) doi:10.1371/journal.pbio.3001310.

22. Puttagunta, R. et al. PCAF-dependent epigenetic changes promote axonal regeneration in the central nervous system. Nat Commun 5, 3527 (2014).

23. Park, K. K. et al. Promoting Axon Regeneration in the Adult CNS by Modulation of the PTEN/mTOR Pathway. Science 322, 963–966 (2008).

24. Liu, K. et al. PTEN Deletion Enhances the Regenerative Ability of Adult Corticospinal Neurons. Nat Neurosci 13, 1075–1081 (2010).

25. Menon, A. S. et al. PATZ1 Reinstates a Growth-Permissive Chromatin Landscape in Adult Corticospinal Neurons After Injury. Preprint at 10.1101/2025.05.14.654166 (2025).

26. van der Velde, A. et al. Annotation of chromatin states in 66 complete mouse epigenomes during development. Commun Biol 4, 239 (2021).

27. Gorkin, D. U. et al. An atlas of dynamic chromatin landscapes in mouse fetal development. Nature 583, 744–751 (2020).

28. Lui, J. C., Chen, W., Cheung, C. S. F. & Baron, J. Broad Shifts in Gene Expression during Early Postnatal Life Are Associated with Shifts in Histone Methylation Patterns. PLOS ONE 9, e86957 (2014).

29. Finn, E. H. & Misteli, T. Molecular basis and biological function of variability in spatial genome organization. Science 365, eaaw9498 (2019).

30. Krishna, N., Menon, A. S., Kumaran, M., Kesireddy, D. K. & Venkatesh, I. PATZ1 remodels the nucleosome landscape to promote chromatin accessibility in injured neurons. 2025.12.22.695949 Preprint at 10.64898/2025.12.22.695949 (2025).

31. Ing-Simmons, E. et al. Spatial enhancer clustering and regulation of enhancer-proximal genes by cohesin. Genome Res 25, 504–513 (2015).

32. Lieberman-Aiden, E. et al. Comprehensive Mapping of Long-Range Interactions Reveals Folding Principles of the Human Genome. Science 326, 289–293 (2009).

33. Li, Y., Hu, M. & Shen, Y. Gene regulation in the 3D genome. Hum Mol Genet 27, R228–R233 (2018).

34. Vermunt, M. W., Zhang, D. & Blobel, G. A. The interdependence of gene-regulatory elements and the 3D genome. J Cell Biol 218, 12–26 (2019).

35. Bouwman, B. A. & de Laat, W. Getting the genome in shape: the formation of loops, domains and compartments. Genome Biol 16, 154 (2015).

36. Dekker, J. & Mirny, L. The 3D Genome as Moderator of Chromosomal Communication. Cell 164, 1110–1121 (2016).

37. Bonev, B. & Cavalli, G. Organization and function of the 3D genome. Nat Rev Genet 17, 661–678 (2016).

38. Rowley, M. J. & Corces, V. G. Organizational principles of 3D genome architecture. Nat Rev Genet 19, 789–800 (2018).

39. Fujita, Y., Pather, S. R., Ming, G. & Song, H. 3D spatial genome organization in the nervous system: From development and plasticity to disease. Neuron 110, 2902–2915 (2022).

40. Dixon, J. R. et al. Chromatin architecture reorganization during stem cell differentiation. Nature 518, 331–336 (2015).

41. Harris, H. L. et al. Chromatin alternates between A and B compartments at kilobase scale for subgenic organization. Nat Commun 14, 3303 (2023).

42. Dixon, J. R. et al. Topological domains in mammalian genomes identified by analysis of chromatin interactions. Nature 485, 376–380 (2012).

43. Nora, E. P. et al. Spatial partitioning of the regulatory landscape of the X-inactivation centre. Nature 485, 381–385 (2012).

44. Sexton, T. et al. Three-dimensional folding and functional organization principles of the Drosophila genome. Cell 148, 458–472 (2012).

45. Phillips-Cremins, J. E. et al. Architectural protein subclasses shape 3D organization of genomes during lineage commitment. Cell 153, 1281–1295 (2013).

46. Rao, S. S. P. et al. A 3D Map of the Human Genome at Kilobase Resolution Reveals Principles of Chromatin Looping. Cell 159, 1665–1680 (2014).

47. Fudenberg, G. et al. Formation of Chromosomal Domains by Loop Extrusion. Cell Reports 15, 2038–2049 (2016).

48. Ray-Jones, H. & Spivakov, M. Transcriptional enhancers and their communication with gene promoters. Cell Mol Life Sci 78, 6453–6485 (2021).

49. Dekker, J., Rippe, K., Dekker, M. & Kleckner, N. Capturing chromosome conformation. Science 295, 1306–1311 (2002).

50. Ilaria Palmisano et al. Three-dimensional chromatin mapping of sensory neurons reveals that promoter-enhancer looping is required for axonal regeneration. Proceedings of the National Academy of Sciences of the United States of America 10.1073/pnas.2402518121 (2024) doi:10.1073/pnas.2402518121.

51. Bonev, B. et al. Multiscale 3D Genome Rewiring during Mouse Neural Development. Cell 171, 557–572.e24 (2017).

52. Hug, C. B. & Vaquerizas, J. M. The Birth of the 3D Genome during Early Embryonic Development. Trends Genet 34, 903–914 (2018).

53. Zheng, H. & Xie, W. The role of 3D genome organization in development and cell differentiation. Nat Rev Mol Cell Biol 20, 535–550 (2019).

54. Blackmore, M. G. et al. Krüppel-like Factor 7 engineered for transcriptional activation promotes axon regeneration in the adult corticospinal tract. Proc. Natl. Acad. Sci. U.S.A. 109, 7517–7522 (2012).

55. Wang, Z. et al. Single-nuclei sequencing reveals a robust corticospinal response to nearby axotomy but overall insensitivity to spinal injury. J. Neurosci. e1508242024 (2025) doi:10.1523/JNEUROSCI.1508-24.2024.

56. Rhodes, C. T. et al. An epigenome atlas of neural progenitors within the embryonic mouse forebrain. Nat Commun 13, 4196 (2022).

57. Tan, L. et al. Changes in genome architecture and transcriptional dynamics progress independently of sensory experience during post-natal brain development. Cell 184, 741–758.e17 (2021).

58. Fraser, J. et al. Hierarchical folding and reorganization of chromosomes are linked to transcriptional changes in cellular differentiation. Mol Syst Biol 11, 852 (2015).

59. Tan, L. et al. Lifelong restructuring of 3D genome architecture in cerebellar granule cells. Science 10.1126/science.adh3253 (2023) doi:10.1126/science.adh3253.

60. Liu, S. et al. Cell type–specific 3D-genome organization and transcription regulation in the brain. Sci Adv 11, eadv2067.

61. Norrie, J. L. et al. Nucleome Dynamics during Retinal Development. Neuron 104, 512–528.e11 (2019).

62. Chen, Z. et al. Increased enhancer-promoter interactions during developmental enhancer activation in mammals. Nat Genet 56, 675–685 (2024).

63. Ghavi-Helm, Y. et al. Enhancer loops appear stable during development and are associated with paused polymerase. Nature 512, 96–100 (2014).

64. Pollex, T. et al. Enhancer-promoter interactions become more instructive in the transition from cell-fate specification to tissue differentiation. Nat Genet 56, 686–696 (2024).

65. Dileep, V. et al. Neuronal DNA double-strand breaks lead to genome structural variations and 3D genome disruption in neurodegeneration. Cell 186, 4404–4421.e20 (2023).

66. Beagan, J. A. et al. Three-dimensional genome restructuring across timescales of activity-induced neuronal gene expression. Nat Neurosci 23, 707–717 (2020).

67. Won, H. et al. Chromosome conformation elucidates regulatory relationships in developing human brain. Nature 538, 523–527 (2016).

68. Fulco, C. P. et al. Activity-by-contact model of enhancer-promoter regulation from thousands of CRISPR perturbations. Nat Genet 51, 1664–1669 (2019).

69. Onrust-van Schoonhoven, A., et al. 3D chromatin reprogramming primes human memory TH2 cells for rapid recall and pathogenic dysfunction. Sci Immunol 8, eadg3917 (2023).

70. Pollex, T. et al. Chromatin gene-gene loops support the cross-regulation of genes with related function. Molecular Cell 84, 822–838.e8 (2024).

71. Marco, A. et al. Mapping the epigenomic and transcriptomic interplay during memory formation and recall in the hippocampal engram ensemble. Nat Neurosci 23, 1606–1617 (2020).

72. Kim, K. et al. Epigenetic memory in induced pluripotent stem cells. Nature 467, 285–290 (2010).

73. Onrust-van Schoonhoven, A., et al. 3D chromatin reprogramming primes human memory TH2 cells for rapid recall and pathogenic dysfunction. Sci Immunol 8, eadg3917 (2023).

74. Kim, T.-K. et al. Widespread transcription at neuronal activity-regulated enhancers. Nature 465, 182–187 (2010).

75. N, Y., et al. Histone H1 loss drives lymphoma by disrupting 3D chromatin architecture. Nature 589, (2021).

76. He, M. et al. Reprogramming of 3D genome structure underlying HSPC development in zebrafish. Stem Cell Res Ther 15, 172 (2024).

77. Plaza-Jennings, A. L. et al. HIV integration in the human brain is linked to microglial activation and 3D genome remodeling. Mol Cell 82, 4647–4663.e8 (2022).

78. Hao, N., Shearwin, K. E. & Dodd, I. B. Programmable DNA looping using engineered bivalent dCas9 complexes. Nat Commun 8, 1628 (2017).

79. Morgan, S. L. et al. CRISPR-Mediated Reorganization of Chromatin Loop Structure. J Vis Exp 57457 (2018) doi:10.3791/57457.

80. Guo, Y. et al. CRISPR Inversion of CTCF Sites Alters Genome Topology and Enhancer/Promoter Function. Cell 162, 900–910 (2015).

81. Wang, H. et al. CRISPR-Mediated Programmable 3D Genome Positioning and Nuclear Organization. Cell 175, 1405–1417.e14 (2018).

82. Schmitt, A. D., Hu, M. & Ren, B. Genome-wide mapping and analysis of chromosome architecture. Nat Rev Mol Cell Biol 17, 743–755 (2016).

83. Chen, S., Zhou, Y., Chen, Y. & Gu, J. fastp: an ultra-fast all-in-one FASTQ preprocessor. Bioinformatics 34, i884–i890 (2018).

84. Heinz, S. et al. Simple combinations of lineage-determining transcription factors prime cis-regulatory elements required for macrophage and B cell identities. Mol Cell 38, 576–589 (2010).

85. Langmead, B. & Salzberg, S. L. Fast gapped-read alignment with Bowtie 2. Nat Methods 9, 357–359 (2012).

86. Open2C et al. Cooltools: Enabling high-resolution Hi-C analysis in Python. PLoS Comput Biol 20, e1012067 (2024).

87. Open2C et al. Bioframe: operations on genomic intervals in Pandas dataframes. Bioinformatics 40, btae088 (2024).

88. Hunter, J. D. Matplotlib: A 2D Graphics Environment. Computing in Science & Engineering 9, 90–95 (2007).

89. Abdennur, N. & Mirny, L. A. Cooler: scalable storage for Hi-C data and other genomically labeled arrays. Bioinformatics 36, 311–316 (2020).

90. Virtanen, P. et al. SciPy 1.0: fundamental algorithms for scientific computing in Python. Nat Methods 17, 261–272 (2020).

91. Dessau, R. B. & Pipper, C. B. [’’R“--project for statistical computing]. Ugeskr Laeger 170, 328–330 (2008).

92. Ito, K. & Murphy, D. Application of ggplot2 to Pharmacometric Graphics. CPT Pharmacometrics Syst Pharmacol 2, e79 (2013).

93. Wolff, J. et al. Galaxy HiCExplorer 3: a web server for reproducible Hi-C, capture Hi-C and single-cell Hi-C data analysis, quality control and visualization. Nucleic Acids Res 48, W177–W184 (2020).

94. Kramer, N. E. et al. Plotgardener: cultivating precise multi-panel figures in R. Bioinformatics 38, 2042–2045 (2022).

