## Supplementary Figure S1 to S6 for "Rewiring of the three-dimensional genome encodes regenerative potential in the adult central nervous system"

Supplementary Figures

Anisha S Menon et al.,

#### Supplementary Figure S1

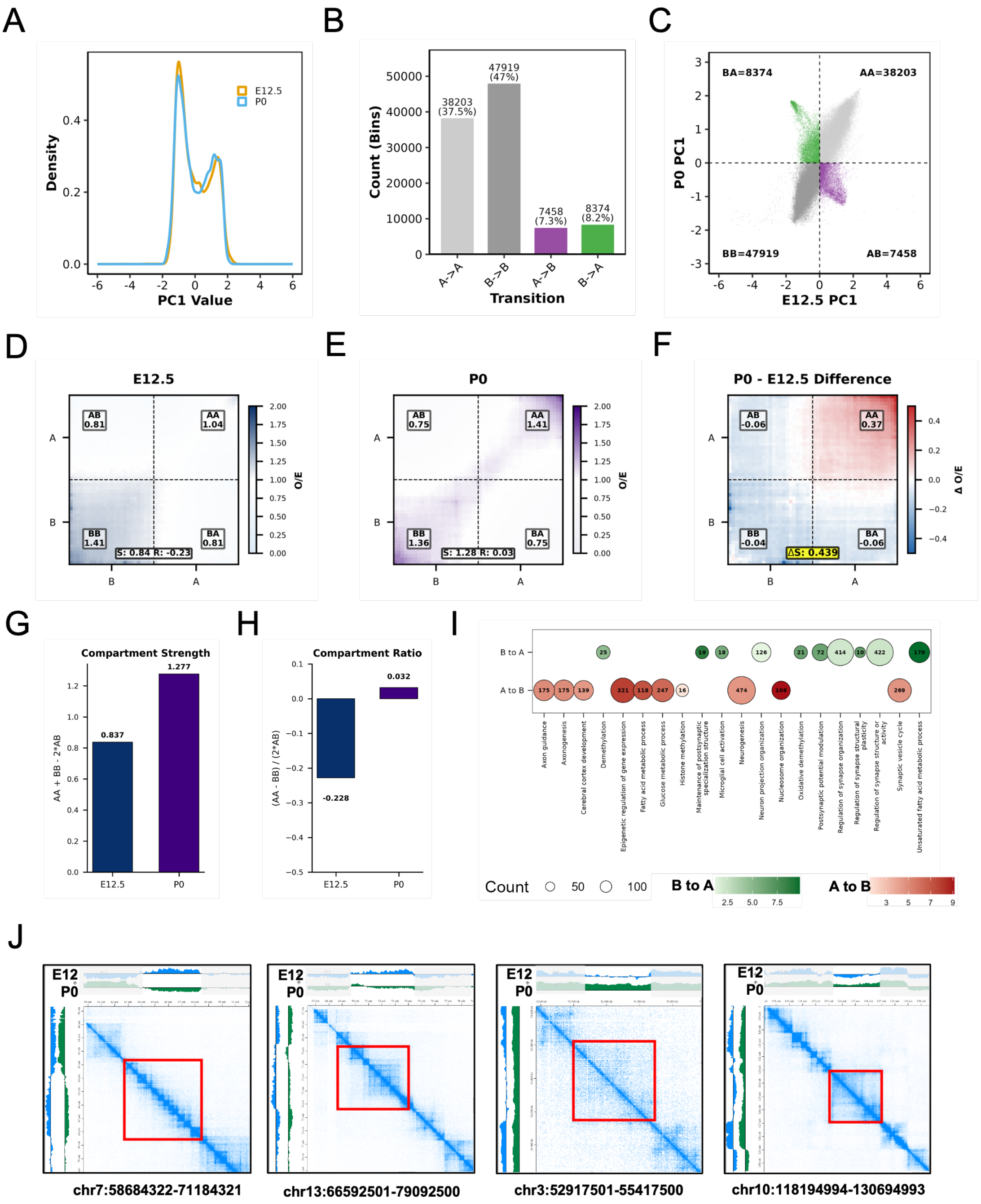


**Supplementary Figure S1. Global Reorganization of Chromatin Compartments During E12.5 to P0 Development.** (A) Distribution of Compartment Scores. Density plot showing the distribution of Principal Component 1 (PC1) values derived from Hi-C data at E12.5 (orange) and P0 (blue). The bimodal distribution reflects genome-wide segregation into transcriptionally active (A) and inactive (B) compartments. (B) Genome-wide Compartment Transitions. Bar chart quantifying the percentage of the genome maintaining or shifting compartment identity between E12.5 and P0. The majority of the genome remains stable (A-to-A and B-to-B), while approximately 16% undergoes A-to-B or B-to-A transitions. (C) Correlation of Compartment

Dynamics. Scatter plot comparing PC1 values at E12.5 versus P0 across the genome (Pearson R = 0.791). Highlighted regions indicate significant compartment transitions: B-to-A (green; n = 6,515) and A-to-B (purple; n = 6,405), representing regions of chromatin activation and repression, respectively. (D) Saddle Plot at E12.5. Observed/expected interaction frequency saddle plot at E12.5. The compartment strength score is 0.84 (blue). The red line demarcates the boundary between A and B compartments. (E) Saddle Plot at P0. Observed/expected interaction frequency saddle plot at P0. The compartment strength score is 1.28 (purple). The red line demarcates the boundary between A and B compartments. (F) Saddle Plot Difference (E12.5 − P0). Differential saddle plot comparing E12.5 and P0, with a difference score of 0.438, reflecting the overall gain in compartmentalization between the two developmental timepoints. (G) Compartment Strength. Bar plot showing compartment strength at E12.5 (0.837) and P0 (1.277), calculated as AA + BB − (2 × AB), where AA, BB, and AB represent the mean observed/expected interaction frequencies within and between compartments. (H) Compartment Ratio. Bar plot showing the compartment ratio at E12.5 (−0.228) and P0 (0.032), calculated as (AA − BB) / (2 × AB). This metric reflects the relative balance of active versus inactive compartment interactions across developmental stages. (I) Gene Set Enrichment Analysis (GSEA). Dot plots representing GSEA results for genes within compartment-switching regions. Genes in regions transitioning from A-to-B between E12.5 and P0 (repressed at P0) are enriched for hematopoietic and muscle development pathways, while genes in regions transitioning from B-to-A (activated at P0) are enriched for synaptic plasticity and neuronal regulation pathways. (J) Visualization of Chromatin Contact Dynamics. Representative Hi-C contact maps for selected genomic loci on chromosomes 3, 7, 10, and 13. Heatmaps contrast contact intensities at E12.5 (blue) and P0 (green), with red boxes highlighting regions of compartment switching or strengthening.

#### Supplementary Figure S2

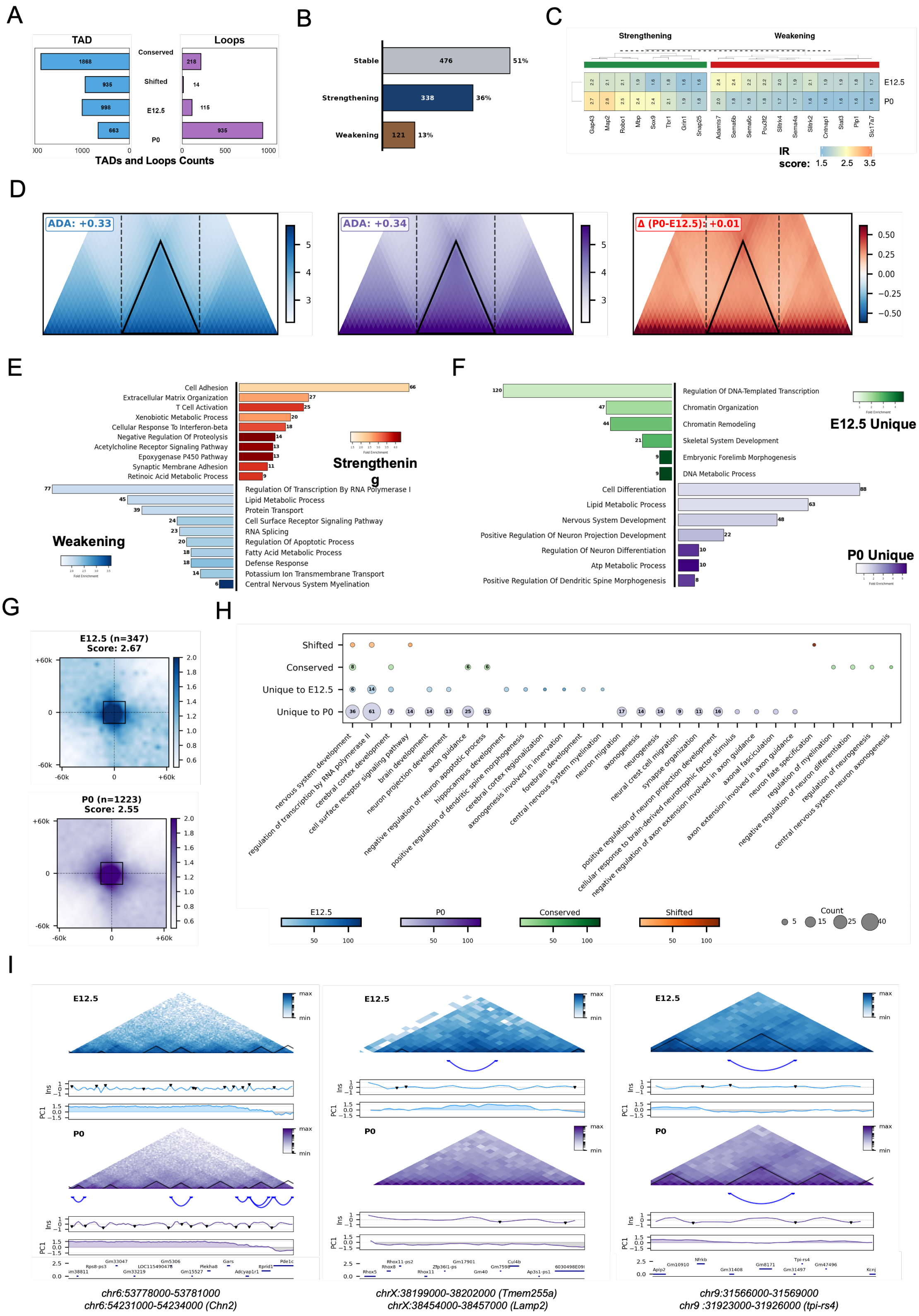

**Supplementary Figure S2. Global Reorganization of Topologically Associating Domains (TADs) and Chromatin Loops During E12.5 to P0 maturation.** (A) Overview of TAD and Loop Dynamics. Bidirectional bar plot summarizing the classification of TADs (left, blue) and loops (right, purple) between E12.5 and P0. Categories include conserved, shifted, unique to

E12.5, and unique to P0 elements, illustrating the extent of structural reorganization across development. (B) TAD Boundary Dynamics. Bar plot quantifying TAD behavior between E12.5 and P0. The majority of TADs are stable (51%), followed by strengthening (36%), and weakening (13%) TADs, highlighting widespread remodeling of domain architecture during this developmental transition. (C) Top Strengthening and Weakening TADs. Heatmap displaying the top 20 TADs ranked by degree of strengthening or weakening between E12.5 and P0, providing a locus-specific view of the most dynamically reorganized chromatin domains. (D) Aggregate Domain Analysis (ADA). ADA plot comparing average TAD insulation at E12.5 (blue; score = 0.33) and P0 (purple; score = 0.34). The modest difference ($\Delta$P0 − E12.5 = 0.01) suggests overall TAD insulation is largely preserved, with subtle global strengthening at P0. (E) Gene Ontology (GO) Enrichment of Dynamic TADs. Bar plot showing the top GO pathways enriched in genes within strengthening TADs (orange) and weakening TADs (blue), linking chromatin domain dynamics to specific biological processes during development. (F) Gene Ontology (GO) Enrichment of Stage-Specific TADs. Bar plot displaying the top enriched GO pathways for genes within TADs unique to E12.5 (green) and unique to P0 (purple), highlighting stage-specific regulatory programs associated with gained or lost domain structures. (G) Aggregate Peak Analysis (APA). APA plot comparing chromatin loop strength at E12.5 (blue; score = 2.67) and P0 (purple; score = 2.55), indicating a modest global reduction in loop intensity from embryonic to postnatal stages. (H) Gene Ontology (GO) Enrichment of Loop Categories. GO enrichment analysis for genes associated with shifted loops, conserved loops, loops specific to P0, and loops specific to E12.5, revealing the biological pathways linked to differential loop organization across developmental stages. (I) Representative Genomic Loci. Genome browser screenshots of selected regions illustrating examples of TAD and loop reorganization between E12.5 and P0, with Hi-C contact maps and corresponding annotations highlighting structural changes at specific loci.

### Supplementary Figure S3

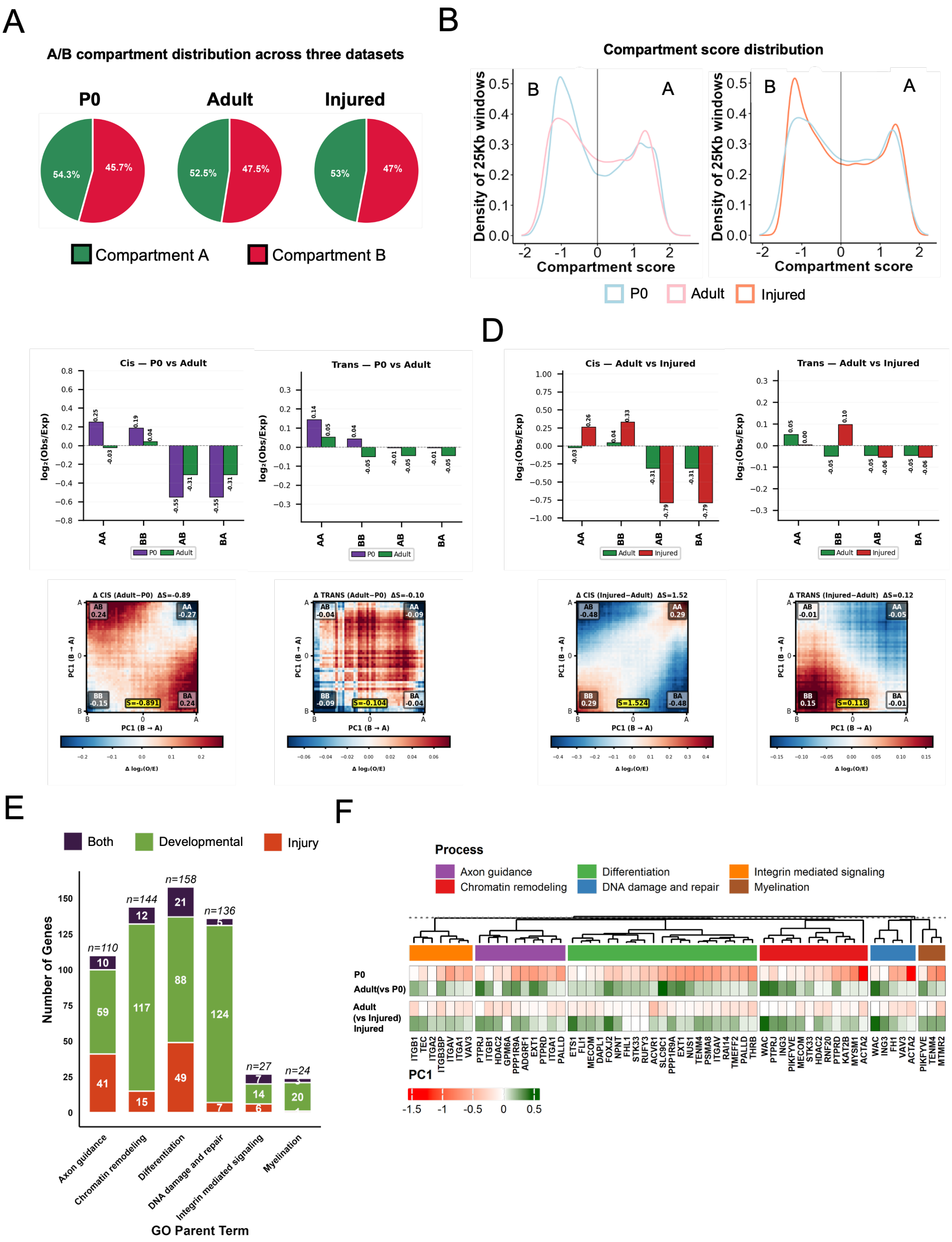


**Supplementary Figure S3: Global Reorganization of Chromatin Compartments Between P0, Adult and Injured Mice** (A) Fraction of genomic regions assigned to A and B compartments across developmental and injury conditions. Pie charts show the proportion of compartments, A (active) and B (inactive), during development and injury conditions. Bar plots compare compartment fractions between Adult and Injured samples. B. Kernel density distributions of compartment scores (PC1) calculated over 25 kb genomic windows for P0, Adult, and Injured datasets illustrates the global strength of compartment segregation. Negative PC1 values correspond to B compartments, while positive PC1 values correspond to A compartments. The vertical line indicates PC1 = 0, marking the boundary between A and B compartments. It illustrates the global strength of compartment segregation. (C) Compartment interaction analysis ( cis and trans) between P0 and adult. Bar charts (top) show the $\log_2$ observed/expected (O/E) interaction frequencies for AA, BB, AB, and BA compartment pairs in cis (left) and trans (right) interactions. Saddle plots (bottom) display the genome-wide difference in compartment interactions ($\Delta \log_2$ O/E) between P0 and Adult, with the segregation strength ($\Delta S$) indicated for cis ($\Delta S = 0.89$) and trans ($\Delta S = 0.10$) interactions. Positive values (red) indicate gained interactions and negative values (blue) indicate lost interactions in adult relative to P0, reflecting a strengthening of compartment segregation during development.(D) Compartment interaction analysis between Adult and Injured conditions. Bar charts (top) show $\log_2$ O/E interaction frequencies for AA, BB, AB, and BA compartment pairs in cis (left) and trans (right). Saddle plots (bottom) display the difference in compartment interactions ($\Delta \log_2$ O/E) between adult and injured, with segregation strength indicated for cis ($\Delta S = -1.52$) and trans ($\Delta S = -0.12$). The negative $\Delta S$ values and the predominance of red in inter-compartment (AB/BA) interactions indicate a loss of compartment segregation following injury, suggesting a shift toward a less defined chromatin compartment structure. (E) Stacked bar chart showing the number of genes associated with shared GO biological process parent terms identified in both the developmental (P0 vs. adult) and injury (adult vs. injured) compartment comparisons. Genes are categorized as developmental only (green), injury only (orange), or common to both comparisons (purple). Total gene counts per term are indicated above each bar, highlighting the degree of gene overlap across biological processes including axon guidance, chromatin remodeling, differentiation, DNA damage and repair, integrin-mediated signaling, and myelination.(F) Heatmap displaying PC1 values for genes shared across developmental and injury GO terms, hierarchically clustered by similarity of compartment identity changes. Rows represent each condition (P0, adult vs. P0, adult vs. injured, and injured) and columns represent individual genes, colored by biological process. PC1 values reflect compartment identity, where positive values (green) indicate active (A) compartment association and negative values (red) indicate inactive (B) compartment association. Genes predominantly reside in the active compartment at P0, shift toward the inactive compartment in adult, and return to a more active compartment state following injury, suggesting a partial reversion to an immature chromatin compartment configuration.

### Supplementary Figure S4

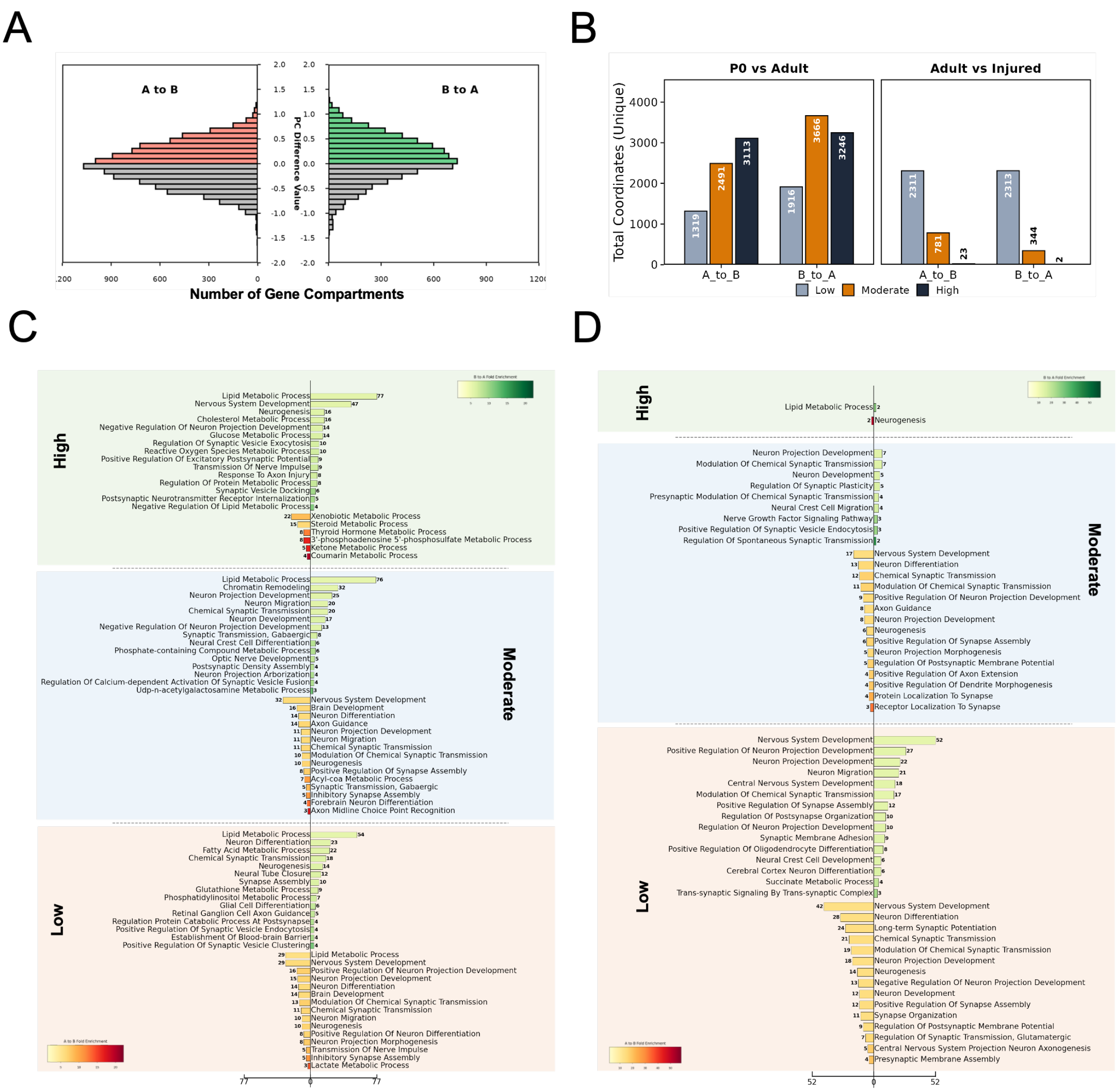


\

**Supplementary Figure S4: Global Gene Ontology analysis of compartment shifts across maturation and injury states.** (A) Distribution of compartment shifts between P0 and Adult states. Histogram representing the frequency of gene compartment transitions. Salmon-colored bars indicate shifts from Compartment A to B, while light green bars represent shifts from Compartment B to A. The grey bars indicate the degree of PC difference value across the number of gene compartments. (B) Quantification of unique compartment coordinates by shift magnitude. Bar plots showing the total number of unique coordinates categorized into Low, Moderate, and High shift bins. Data is presented for two comparative states: P0 vs. Adult (left) and Adult vs. Injured (right). Note the significant decrease in high-magnitude shifts in the Adult vs. Injured dataset compared to the developmental P0 vs. Adult dataset. (C) Gene Ontology (GO) Enrichment for P0 vs. Adult transitions. Enrichment analysis of biological processes categorized by shift magnitude (High, Moderate, and Low). The green gradient bars indicate enrichment for genes shifting from B to A, while red gradient bars indicate enrichment for genes shifting from A to B. Terms include Lipid Metabolic Process,Nervous System Development and Synaptic Transmission. (D) Gene Ontology (GO) Enrichment for Adult vs. Injured transitions. GO analysis for the Adult vs. Injured dataset following the same magnitude categorization (High, Moderate, and Low). Similar to panel C, the green bars represent B to A shifts and red bars represent A to B shifts. This panel highlights the functional genomic response to injury, showing a distinct profile compared to developmental shifts.

#### Supplementary Figure S5

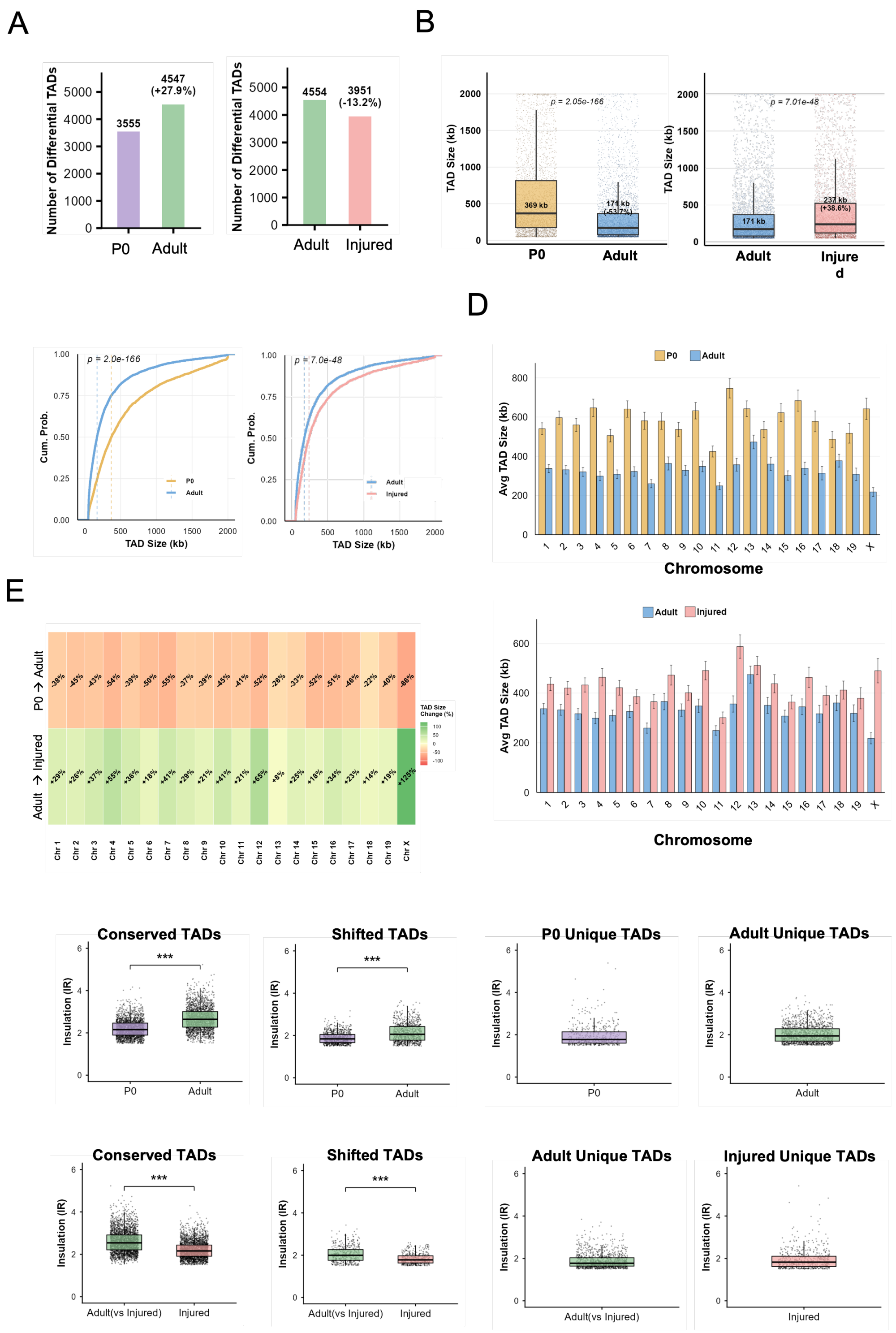

**Supplementary Figure S5: TAD analysis shows significant changes in TAD sizes between P0, Adult and Injured Mice** (A) The bar charts display the change in the number of differential TADs across development and injury. (Left) The number of TADs increased by 27.9% during development from P0 to Adult. (Right) The number of TADs decreased by 13.2% upon injury. (B) Box plots showing global shifts in TAD size (kb). (Left) TADs significantly contract during development (P0 vs. Adult), with the median size decreasing from 369 kb to 171 kb (-53.7%; $p < 2.1e-166$, Wilcoxon rank-sum test ). (Right) TADs significantly expand following injury, with the median size increasing from 171 kb to 237 kb (+38.6%; $p < 7.1e-48$, Wilcoxon rank-sum test ). C. Empirical Cumulative Distribution Function (ECDF) plot of TAD sizes during development (P0 vs. Adult) and Injury ( Adult vs Injured). During development Adult TADs (blue) are significantly smaller than P0 TADs (yellow) ($p = 2.05 \times 10^{-166}$, Wilcoxon rank-sum test). Injury causes a significant shift toward larger TAD sizes (red curve shifted right), ($p = 7.01 \times 10^{-48}$, Wilcoxon rank-sum test). (D) The figure shows chromosome wise TAD size distribution during development and upon injury. (E) Related to Fig. D. The heatmap quantifies percentage TAD size change chromosome wise. (F, G) The box plots represent the distribution insulation score of TADs categorized into Conserved, Shifted, and Unique TADs (p-value < 0.001, Mann Whiteny Unpaired Test)

### Supplementary Figure S6

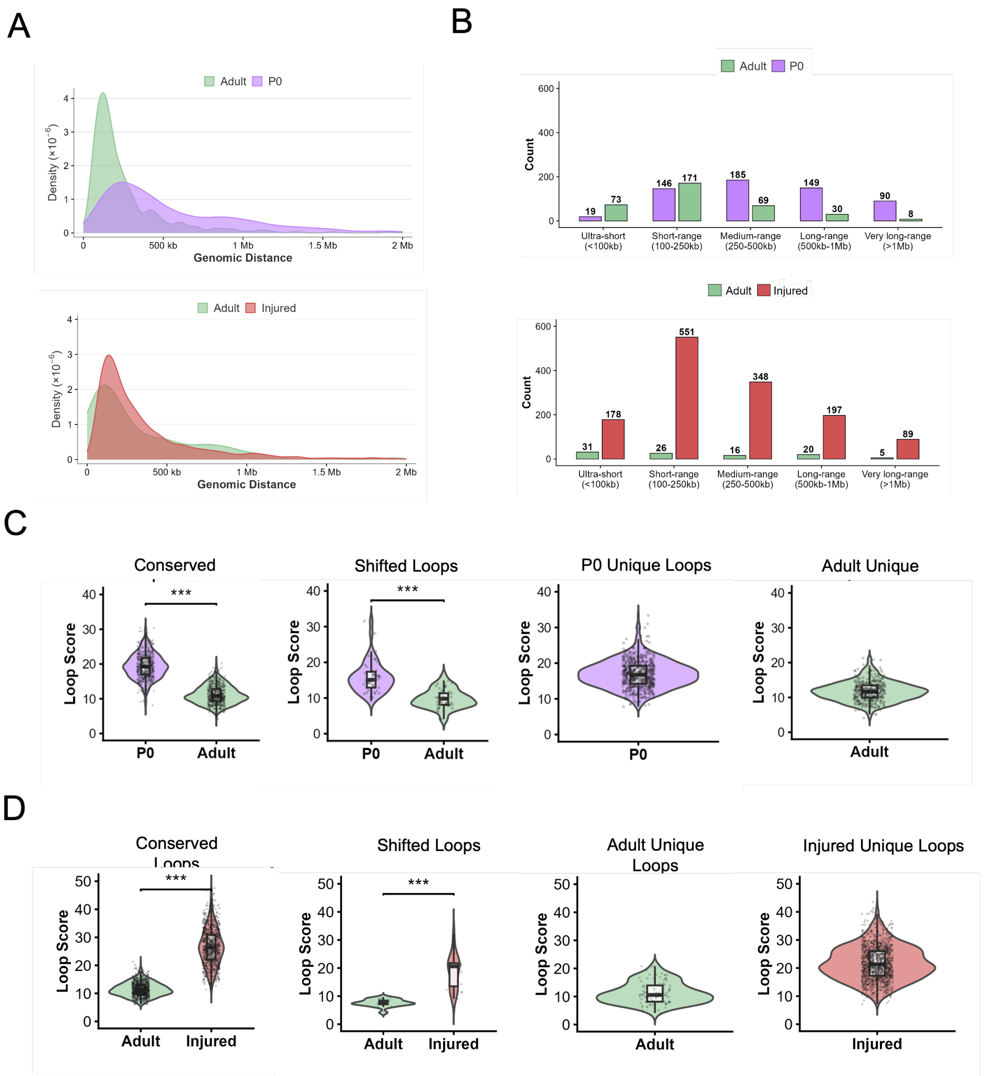


**Supplementary Figure S6: Characterization of loop dynamics across development and injury.** (A) Kernal Density Enrichment plot of Hi-C chromatin loop sizes. (Top) Distribution of anchor-to-anchor distances in Adult (green) versus P0 (purple) samples, highlighting a distinct peak of shorter loops in adults compared to broader, longer-range interactions in P0. (Bottom) Comparison of loop sizes in Adult Uninjured (dark green) versus Injured (red) conditions, showing a pronounced shift toward shorter genomic distances following injury. *(For both panels, the x-axis indicates genomic distance up to 2 Mb; the y-axis indicates probability density × $10^{-6}$). (*B) Quantification of chromatin loops by size category. (Top) Bar plot comparing P0 (purple) and Adult (green) stages. P0 exhibits higher counts of medium-to-long range loops, while adults shift toward shorter interactions (<250kb). (Bottom) Comparison of Adult Uninjured (green) and Injured (red) conditions, demonstrating a massive overall increase in loop counts following injury, particularly in the 100–500kb range. (C and D) Loop score distributions across four loop categories - Conserved, Shifted, and unique loops to each dataset. Violins show the full distribution, internal boxplots indicate median and interquartile range, and grey dots represent individual loops. Statistical comparisons performed using the Wilcoxon rank-sum test (*** $p < 0.001$, **** $p < 0.0001$)
